# Extremely oligotrophic and complex carbon degrading microaerobic bacteria from Arabian Sea oxygen minimum zone sediments

**DOI:** 10.1101/2023.10.31.564988

**Authors:** Jagannath Sarkar, Mahamadul Mondal, Sabyasachi Bhattacharya, Subhajit Dutta, Sumit Chatterjee, Nibendu Mondal, Saran N, Aditya Peketi, Aninda Mazumdar, Wriddhiman Ghosh

## Abstract

Sediments underlying marine hypoxic zones are huge sinks of unreacted complex organic matter, where despite acute O_2_-limitation obligately aerobic bacteria thrive, and steady depletion of organic carbon takes place within a few meters below the seafloor. However, little knowledge exists about the sustenance and complex carbon degradation potentials of aerobic chemoorganotrophs in these sulfidic ecosystems. We isolated and characterized a number of aerobic bacterial chemoorganoheterotrophs from across a ∼3 m sediment horizon underlying the perennial hypoxic zone of the eastern Arabian Sea. High levels of sequence correspondence between the isolates’ genomes and the habitat’s metagenomes and metatranscriptomes illustrated that the strains were widespread and active across the sediment-cores explored. The isolates catabolized several complex organic compounds of marine and terrestrial origins in the presence of high or low, but not zero, O_2_. Some of them could also grow anaerobically on yeast extract or acetate by reducing nitrate and/or nitrite. Fermentation did not support growth, but enabled all the strains to maintain a fraction of their cell populations over prolonged anoxia. Under extreme oligotrophy, limited growth followed by protracted stationary phase was observed for all the isolates at low cell density, amid high or low, but not zero, O_2_ concentration. While population control and maintenance could be particularly useful for the strains’ survival in the critically carbon-depleted layers below the explored sediment-depths (core-bottom organic carbon: 0.5-1.0 % w/w), metagenomic data suggested that *in situ* anoxia could be surmounted via potential supplies of cryptic O_2_ from previously-reported sources such as *Nitrosopumilus* species.

## INTRODUCTION

Marine sediments constitute one of the largest microbiomes on Earth (Hoshino et al. 2020), and act as the largest sink of organic carbon within the biosphere (Kallmeyer et al. 2012; Jørgensen et al. 2022; Ruben et al. 2023). Particularly on continental shelves and slopes, productivity in the oceanic water column, rate of sedimentation, and the extent of organic matter delivery to the seafloor are all very high; subsequently O_2_, followed by the less energy-efficient respiratory substrates such as nitrate, and oxidized ions of manganese and iron, gets depleted within relatively shallow sediment-depths (Canfield 1994; Seiter et al. 2005; Burdige 2007; Komada et al. 2016; Bhattacharya et al. 2021; Sarkar et al. 2022). This phenomenon is further accentuated in the mid-oceanic oxygen minimum zones (OMZs) that occur perennially along the western margins of the Earth’s continents, and are considered to be the crucial bellwethers of ocean warming, deoxygenation, and acidification caused by global climate change (Wright et al. 2012; Long et al. 2021).

In the northwestern extension of the Indian Ocean a vast body of hypoxic water extends across the Arabian Sea and impinges upon a large shelf/slope area along the continental margins of India, Oman, Pakistan, Yemen, and Cape Guardafui in extreme north-eastern Somalia (Wyrtki 1971; Bange et al. 2005; Naqvi et al. 2006, Banse et al. 2014). This marine territory having dissolved O_2_ concentration perennially below 20_µM is regarded as the thickest and largest of all the oxygen minimum zones (OMZs) of the global ocean (Acharya and Panigrahi 2016; Long et al. 2021). Its upper boundary is positioned between 100 and 150 meters below the sea level (mbsl), the lower boundary between 1000 and 1200 mbsl, and the underlain benthic area (∼230,440 km^2^) constitutes more than a quarter of the Earth’s total (∼764,000 km^2^) naturally-hypoxic seafloor (Helly and Levin 2004; Acharya and Panigrahi 2016). As photosynthetic biomass from the highly productive euphotic zone descends down the water column, rapid biodegradation takes place, causing intense oxygen depletion and subsequent denitrification in the 100-1200 mbsl zone (Jayakumar et al. 2004; Lam and Kuypers 2011). However, due to hypoxic conditions in the water column, a large fraction of the complex organic matter evades degradation while sinking, and settles on the seafloor in its unreacted and labile form (Van Mooy et al. 2002; Cavan et al. 2017; Jessen et al. 2017). The deposition and degradation dynamics, i.e. the concentration, chemical nature, and fate of the organic matter that is delivered to the seafloor, determine the microbiome structure and function in the sediment system across the continental margins (Bhattacharya et al. 2021). With increasing water-depth and distance from the coast, sedimentation rate decreases, and the organic matter delivered to the seafloor gets exposed to O_2_ for longer periods of time. As a consequence, irrespective of the O_2_ concentration of the bottom-water, faster breakdown of complex organic matter into simpler compounds takes place in the sediment surface (Canfield 1994; Kristensen et al. 1995; Burdige 2007; Middelburg 2019; Bhattacharya et al. 2021). Across the sediment-depths, this leads to an overall intensification of the simple-fatty-acids-requiring anaerobic metabolisms such as sulfate reduction and methanogenesis (Fernandes et al. 2018; Bhattacharya et al. 2021).

Whereas post-burial degradation of complex organic matter in the top-most strata of these sulfide-containing sediment systems is attributable to aerobic bacterial catabolism (Bhattacharya et al. 2021), it is remarkable that obligately aerobic bacteria are present and active (Bhattacharya et al. 2020), alongside sulfate-respiring anaerobes (Fernandes et al. 2018), across a ∼3 meter sediment horizon of the Arabian Sea OMZ (ASOMZ), i.e. from the seafloor up to the depths where both organic carbon and sulfate are depleted and methane biogenesis begins (Fernandes et al. 2018). The presence of metabolically active and potentially growing aerobic bacterial communities up to the sulfate-methane transition zone of this sediment system is intriguing because there is little biogeochemical feasibility of O_2_ production *in situ*, nor there exists any apparent chance of O_2_ influx from elsewhere (Bhattacharya et al. 2019). However, notwithstanding their ecological idiosyncrasy, the aerobic bacteria of ASOMZ sediments warrant an investigation for any potential role(s) that they might have in the degradation of the complex organic carbon which is abundant in the ecosystem and fuels organoclastic sulfate reduction *in situ* (Fernandes et al. 2018). Furthermore, their nutritional capabilities and physiological adaptations need to be examined in the context of the progressive scarcity of oxidants as well as carbon and energy, which characterizes their habitat. Thus, on one hand, it is necessary to have pure culture isolates of aerobic bacteria from ASOMZ sediments and assess them for their respiratory potentials, complex carbon degradation capabilities, and oligotrophy; on the other hand, it is imperative to re-examine the metagenomes and metatranscriptomes of the habitat, in the light of the latest microbiological literature, for potential source(s) of cryptic biogenic O_2_ that might sustain the aerobic bacteria in this highly-reduced sediment system. Plausible hypotheses put forth on the latter issue might well be tested afterwards by future studies of microbiology and/or geochemistry.

With the abovementioned objectives the current study isolated and characterized aerobic bacterial chemoorganoheterotrophs from across ∼3 meter sediment horizons underlying the heart of the ASOMZ, off the west coast of India (coring stations located at 530 and 580 mbsl water-depths). The strains isolated were investigated for their capabilities of growing under aerobic, microaerobic and anaerobic conditions, via degradation of polymeric carbohydrates, aromatic compounds, simple fatty acids and sugars, and also extremely low absolute concentrations of organic carbon. Population ecology, and *in situ* functionality, of the new isolates were assessed via whole genome sequence analysis, followed by investigations of genome-metagenome and genome-metatranscriptome correspondence. Finally, the metagenomes retrieved at 15–30 cm intervals, along the ∼3 m sediment-cores investigated, were explored for putative signatures of the ammonia-oxidizing *Nitrosopumilus* species that has recently been reported as a potential supplier of biogenic O_2_ in marine ecosystems (Walker et al. 2010; Kraft et al. 2022).

## MATERIALS AND METHODS

### Sample collection

Samples from the ∼3 m long sediment-cores SSK42/5 and SSK42/6 retrieved on board the research cruise RV *Sindhu Sankalp* 42 (SSK42 conducted in December 2012) were used for the present studies of microbial ecology. The two cores, collected from 580 mbsl (latitude: 16°49.88’_N, Longitude: 71°58.55’_E) and 530 mbsl (latitude: 16°50.03’_N, Longitude: 71°59.50’_E) of the eastern Arabian Sea, across the western Indian margin, respectively, were investigated using culture-dependent as well as culture-independent methods at an average interval of 20 cm. For all the sediment-depths explored along the two cores rich in silt and clay, sample replicates were collected for the investigation of pure culture isolates, metagenomes, and metatranscriptomes, as described previously (Fernandes et al. 2018; Bhattacharya et al. 2020, 2021; Mandal et al. 2020).

### Isolation of aerobic chemoorganoheterotrophs

Chemoorganoheterotrophic microbial strains were enriched and isolated in the presence of oxygen from the bulk sediments of SSK42/5 and SSK42/6. For this purpose, sediment samples from three discrete layers were mixed homogeneously for each core but not across the two different cores: 1 g sample each from 0, 140 and 295 centimeters below the seafloor (cmbsf) was taken for SSK42/5, while 1 g sample each from 2, 120 and 250 cmbsf was taken for SSK42/6. For each of the two pooled samples, enrichment and isolation was separately carried out on three different culture media: (i) phosphate buffered artificial seawater (ASW, pH 7.5) medium (Bhattacharya et al. 2020) containing high concentration (0.5 g L^-1^) of yeast extract as the energy and carbon source (this medium has been referred to as ASW_HY here onward); (ii) ASW medium containing low concentration (0.001 g L^-1^) of yeast extract as the energy and carbon source (this medium is referred to as ASW_LY); and (iii) basal bicarbonate buffered (BBB, pH 7.5) medium (Bhattacharya et al. 2020) containing sodium acetate (10 mM) as the energy and carbon source (this medium is referred to as BBB_A).

Using the two core-specific bulk (pooled) sediment samples and three different media types, overall six lines of enrichment and isolation were created. For each of these, 1 g of the relevant bulk sediment sample was added to 100 mL of the applicable culture medium contained in a 250 mL cotton-plugged Erlenmeyer flask having normal air in the headspace. The sediment-medium mixture was incubated aerobically for 24 hours at 15°C (this was the approximate *in situ* temperature), following which the sediment particles were allowed to settle down and the turbid supernatant (having OD_600_ ∼0.5) was diluted serially and spread on agar plates of the same medium in which enrichment had taken place. These plates were incubated aerobically at 15°C, and from the biomass that appeared on them, pure culture strains were isolated after several rounds of dilution streaking. Since all the new isolates could grow on ASW_HY, they were maintained in that medium. The strains were taxonomically classified up to the lowest possible category with the help of methods elaborated elsewhere (Saha et al. 2019); they were also deposited for public accession to the National Centre for Microbial Resource, National Centre for Cell Science, Pune, India.

### Aerobic, microaerobic and anaerobic growth

For an aerobic growth experiment, a seed culture of the test strain was first generated by inoculating 20 ml ASW_HY broth with a loopful of inoculum from the working culture of the strain and then incubating the culture aerobically at 15°C. Cells were harvested from the exponentially growing seed culture, washed twice with and resuspended in 0.9% NaCl, and finally added to the experimental medium in such a way that the initial (0 h) cell density of the test culture came in the range of either 10^6^ or 10^3^ colony forming units (CFUs) mL^-1^.

For an anaerobic or microaerobic growth experiment, seed culture preparation and test culture inoculation were carried out in the same way as above except for the fact that the washing of harvested cell pellets, as well as the inoculation of the test culture, was carried out within an H35 Hypoxystation (Don Whitley Scientific, UK), which was pre-set to 0 or 0.1% partial pressure of O_2_ respectively (in either case humidity was maintained at 75%). Inside the H35, anaerobic condition (anoxia) was created via constant flow of N_2_, CO_2_ and H_2_ at a volumetric ratio of 8:1:1, while microaerobic condition (hypoxia) was created by flowing N_2_, CO_2_ and O_2_ at a volumetric ratio of 9:0.99:0.01. Furthermore, in case of an aerobic or microaerobic growth experiment the culture flask was cotton plugged, whereas for an anaerobic experiment, the culture bottle was screw-capped, and incubated inside the H35 Hypoxystation (in all the cases incubation temperature was 15°C). In this way, all aerobically grown cultures had 21% (v/v) O_2_ in the head-space of the flasks, whereas microaerobic and anaerobic cultures had 0.1% (v/v) O_2_, and zero partial pressure of O_2_, in the head-space of the flasks, respectively.

So far as the culture media were concerned, those used for anaerobic growth experiments were supplemented with the following electron acceptors individually or collectively: (CH_3_)_3_NO (27 mM), (CH_3_)_2_SO (56 mM), Fe_2_O_3_ (125 mM), MnO_2_ (1 mM), NaNO_3_ (4 mM), NaNO_2_ (4 mM), and Na_2_SO_4_ (10 mM). Furthermore, separate anaerobic growth experiments were carried out by supplementing the media with 40 mM Fe_2_O_3_ and 17 g L^-1^ humic acids (Benz et al. 1998; Bhattacharya et al. 2020). Media used for anaerobic or microaerobic experiments contained thioglycolate (0.5 g L^-1^) and resazurin (0.0001_g L^-1^) for the purpose of deoxygenation, and subsequent monitoring of the O_2_ level, respectively. Cultures exhibiting anaerobic growth via nitrate (NO_3_) or nitrite (NO_2_) reduction were measured for the concentration of these ions in the spent media by means of ion chromatography carried out using an Eco IC (Metrohm AG, Switzerland) furnished with a conductivity-based detector (Metrohm IC detector 1.850.9010). Separation of anions was carried out using the anion exchange column Metrosep A Supp5 - 250/4.0 (Metrohm AG). The eluent comprised a mixture of NaHCO_3_ (1.0 mM) and Na_2_CO_3_ (3.2 mM), while the regenerant encompassed sulfuric acid (100 mM); 100 µL of 100-fold diluted sample was injected in every chromatographic run, which in turn involved a flow rate of 0.7 mL min^-1^. Chemicals standards provided by the platform manufacturer (Metrohm AG) were utilized to generate the calibration curves for the quantification of NO_3_ and NO_2_ ions; fluctuation range for each concentration measured was ±0.2 ppm.

Colony forming units (CFUs) present in a given experimental culture at any time-point of its incubation (this included the 0 h of incubation) was counted by taking 1 mL of the culture, diluting it by increasing logs of 10, then plating the different dilution grades in triplicates of ASW_HY agar, and finally counting the colonies appearing therein. In order to obtain the CFU count of 1 mL of the culture, colony counts recorded in the different dilution plates were multiplied by their respective dilution factors, following which the values were added up and averaged across the various dilution grades and replica plates generated. For aerobic growth experiments dilution plating was carried out inside a Class II Biological Safety Cabinet (Esco, Singapore), but in case of microaerobic and anaerobic growth, plating was carried out inside the H35 Hypoxystation; in all the three cases, incubation of the CFU-counting plates was carried out under aerobic condition.

### Testing complex organic compound utilization

Isolates were tested for possible growth via catabolism of different complex organic compounds of marine and terrestrial origin as sole sources of energy and carbon. Tested carbon compounds included agar, alginate, carrageenan and chitosan (in nature, these are mostly of marine origin); cellulose, pectin and xylan (these originate mostly in terrestrial organisms); and benzoate and starch (these are produced by marine as well as terrestrial organisms). ASW medium was amended with one of the abovementioned carbon compounds at a time (as a sole carbon source), and axenic cultures of each isolate were incubated individually in the nine different types of chemoorganoheterotrophic media formulated in the process. Agar (Type-I agar, Himedia, India) and carrageenan (Kappa-carrageenan, Merck, Germany), the two typical cell wall components of the red algae (Rhodophyceae), were added to ASW at the concentrations 2 g L^-1^ and 4 g L^-1^ respectively. Chitosan (Sigma-Aldrich, USA), a derivative of chitin found in the exoskeleton of shellfish, was provided at a concentration of 4 g L^-1^. Alginate (sodium salt of alginate, Himedia), found in the cell wall of marine macroalgae belonging to the family Phaeophyceae (brown algae), was added a concentration of 10 g L^-1^. Cellulose (sodium salt of carboxymethylcellulose, Himedia), the main component of the cell wall of land plants, was provided at a concentration of 5 g L^-1^. Xylan (extracted from beechwood, Himedia), a major component of hemicellulose; and pectin (Himedia), a cell wall polysaccharide of terrestrial plants, were added at a concentration of 10 g L^-1^ each. Benzoate (sodium salt, Himedia) and water-soluble starch (Himedia) were added at the concentrations 5 mM and 4 g L^-1^ respectively.

### Test of oligotrophy and fermentative abilities

ASW medium supplemented with extremely low concentration of yeast extract (1 × 10^-12^ g L^-1^) as the energy and carbon source (ASW_ELY) was used to test the oligotrophic growth potentials of the new isolates, under aerobic, microaerobic, or anaerobic condition, starting with a bacterial density in the range of either 10^3^ or 10^6^ CFU mL^-1^.

Whether the isolates could grow or survive (maintain a fraction of the cell population in viable state) by means of fermentative metabolism was tested in ASW supplemented with pyruvate (5 g L^-1^ sodium pyruvate, Sisco Research Laboratories Pvt. Ltd., India) or glucose (10 g L^-1^ D-glucose, Merck, Germany), under anaerobic condition. Pyruvate, the end product of glycolysis, is an important intermediate of the central carbohydrate metabolism. It is the common substrate utilized by all fermentative bacteria: in some cases it supports growth under anaerobic condition (Cocaign-Bousquet et al. 1996; Wagner et al. 2005), while in others, including in a number of obligate aerobes, it helps preserve cell populations in the absence of O_2_ (Eschbach et al. 2004; Pinchuk et al. 2011).

### Phylogenetic and genomic characterizations of the new isolates

Genomic DNA was prepared from the stationary-phase cultures of the new isolates using PureLink Genomic DNA Mini Kit (Thermo Fisher Scientific, USA). The 16S rRNA gene of each strain was amplified using PCR, and sequenced on a 3500xL Genetic Analyzer (Thermo Fisher Scientific), using the Bacteria-specific universal primers 27f and 1492r (Crocetti et al. 2000). The nearly complete 16S rRNA gene sequence of each strain was analyzed to identify its closest phylogenetic relative as described earlier (Ghosh et al. 2006).

Whole genome shotgun sequencing was carried out for all the isolates using the Ion S5 (Thermo Fisher Scientific) and MinION (Oxford Nanopore Technologies, UK) next generation DNA sequencing technologies. Datasets of raw reads obtained using the two technologies were deposited in the Sequence Read Archive of National Center for Biotechnology Information (NCBI), USA, under the BioProject PRJNA309469, with accession numbers given in Table S1. For each genome, Ion S5 reads having Phred score above 20 and length above 100 nucleotides, and MinION reads having quality value greater than 10, were assembled together using Unicycler v0.5.0 in hybrid assembly format (Wick et al. 2017). The genomes were annotated using the Prokaryotic Genome Annotation Pipeline (PGAP) of NCBI which is located at https://www.ncbi.nlm.nih.gov/genome/annotation_prok/. The catalog of protein-coding sequences (CDSs) or genes obtained from PGAP for each genome was annotated again via search against the eggNOG database v5.0 (Huerta-Cepas et al. 2019) using the online utility eggNOG-mapper v2.1.9 (Cantalapiedra et al. 2021) in default mode. Furthermore, to specifically identify genes encoding different glycoside hydrolases, each PGAP-derived catalog was re-annotated by searching against the dbCAN3 and CAZy databases using HMMER (Zheng et al. 2023) and Diamond (Buchfink et al. 2015) with default parameters respectively. The KEGG (Kyoto Encyclopedia of Genes and Genomes) automatic annotation server (KAAS) was used for ortholog assignment and pathway mapping within the genomes (Moriya et al. 2007).

The annotated complete genome sequences obtained in this way were deposited in the NCBI GenBank, under the BioProject PRJNA309469, with accession numbers mentioned in Table S2 (the assembly statistics have also been presented in the same supplementary table). Furthermore, taxonomic identification of the isolated strains up to the lowest possible taxon was carried out on the basis of their genome-genome relatedness with closest phylogenetic relatives, determined in terms of digital DNA-DNA hybridization (dDDH) values, using Type Strain Genome Server (TYGS; Meier-Kolthoff and Göker 2019) of the Leibniz Institute DSMZ - German Collection of Microorganisms and Cell Cultures GmbH. dDDH value of > 70% with respect to the type of a known species was considered to be the criterion for ascribing a new isolate to that species (Meier-Kolthoff and Göker 2019). Furthermore, using their complete genome sequences, phylogeny of the new isolates, in relation to their closest genomic relatives, was reconstructed on the basis of conserved marker gene sequences, using the Up-to-date Bacterial Core Gene pipeline v3.0 (Na et al. 2018). The best fit nucleotide substitution model was selected using the software ModelTest-NG v0.1.7 (Darriba et al. 2020). Subsequent to this, a maximum likelihood tree was generated using the software RAxML v8.2.12 (Stamatakis 2014). The software called Interactive Tree of Life v6.7.5 (Letunic and Bork, 2019) was used to visualize the phylogram generated in this way. In the reconstructed phylogeny, interpretation of nucleotide substitutions was carried out using the generalized time-reversible model that considers the proportion of invariable sites and/or the rate of variation across the sites.

### Metagenomics

Duplicate metagenomes were prepared from the discrete sediment samples of SSK42/5 and SSK42/6 using PowerSoil DNA Isolation Kit (Mo Bio Laboratories Inc., USA), and sequenced individually with the help of the high-throughput platforms Ion PGM and Ion Proton (Thermo Fisher Scientific, USA), as described elsewhere (Bhattacharya et al. 2020; Mandal et al. 2020). Sediment-depths explored included 0, 15, 45, 60, 90, 120, 140, 160, 190, 220, 260 and 295 cmbsf for SSK42/5 and 2, 30, 45, 60, 75, 90, 120, 135, 175, 220, 250, 265 and 275 cmbsf for SSK42/6. Post merging of the sequence dataset pair obtained for a given sediment sample (data throughput for all the samples given in Table S3), all reads having Phred score ≥ 20 (Q20) and length ≥ 50 nucleotides were subjected to *de novo* contig assembly using Megahit v1.2.9 (Li et al. 2015). While 60% to 94% of available reads participated in the assembly for the different samples, average coverage ranged between 6X to 20X (Table S3). Genes or open reading frames (ORFs), followed by CDSs, were detected and annotated within ≥ 100 nucleotide contigs using Prodigal v2.6.3 (Hyatt et al. 2010), and eggNOG-mapper v2.1.9 (Cantalapiedra et al. 2021) with the help of eggNOG database v5.0 (Huerta-Cepas et al. 2019), respectively.

To determine the level of genome-metagenome correspondence (i.e. the relative abundance of the individual genomes within a metagenome), all quality-filtered metagenomic reads available for each sediment sample (i.e. all Q20 reads taken after merging the duplicate datasets) were mapped onto the bacterial genomes (chromosomes and plasmids taken together) one by one using Bowtie2 v2.4.5 in default mode (Langmead and Salzberg 2012). After knowing what percentage of metagenomic reads from a given sediment sample mapped onto a particular genome, the alignment output file obtained from the Bowtie2 analysis was used as input data for another analysis by the software package BEDTools v2.31.1 (Quinlan and Hall, 2010) to determine what percentage of the total length of the genome in question was covered by the mapped metagenomic reads (this is referred to as coverage breadth).

### Metatranscriptomics

Extraction of metatranscriptomes (using RNA PowerSoil Total RNA Isolation Kit, Mo Bio Laboratories Inc.), removal of rRNAs, and 2 × 150 nucleotides paired-end sequencing on a HiSeq4000 platform (Illumina Inc., USA) were carried out as communicated previously (Bhattacharya et al. 2020, 2021; Mandal et al. 2020). For a given sediment sample, all metatranscriptomic reads having Phred score ≥ 20 were assembled *de novo* using rnaSPAdes v3.15.4 (Bushmanova et al. 2019). ≥ 99.2% reads from each metatranscriptomic dataset analyzed contributed to its assembly rendered at an average coverage of 188X–1120X (Table S4). Genes or ORFs, followed by CDSs, were delineated and annotated within ≥ 100 nucleotide contigs using Prodigal v2.6.3 (Hyatt et al. 2010), and eggNOG-mapper v2.1.9 (Cantalapiedra et al. 2021) with the help of eggNOG database v5.0 (Huerta-Cepas et al. 2019), respectively. In order to specifically identify genes encoding different glycoside hydrolases, each Prodigal-derived CDS-catalog was annotated by searching against the dbCAN3 database using HMMER (Zheng et al. 2023) and the CAZy database using Diamond (Buchfink et al. 2015). All the search engines mentioned above were operated with default parameters.

To determine the level of genome-metatranscriptome correspondence, all quality-filtered, mRNA-specific metatranscriptomic reads available for a given sediment sample were mapped onto the bacterial genomes (chromosomes and plasmids taken together) one by one using Bowtie2 v2.4.5 in default mode. In order to remove potential rRNA-related reads that were there in the metatranscriptomic sequence datasets, the latter were searched with reference to the rRNA gene sequence database SILVA v138.1 (Quast et al. 2013) using the software Bowtie2 v2.4.5 in default mode. After knowing what percentage of metatranscriptomic reads from any sediment sample mapped onto a given genome, coverage breadth (what percentage of that genome’s total length was covered by the mapped metatranscriptomic reads) was determined using BEDTools v2.31.1.

### Detection of *Nitrosopumilus*-specific 16S rRNA gene fragments within the metagenomic DNA samples

The *Nitrosopumilus-*specific primer pair N_pumi_420F (5_-TGTTGAATAAGGGGTGGGCA-3_) and N_pumi_981R (5_-CCACCTCTCAGCTTGTCTGG-3_), obtained using the Primer-BLAST program available online (Ye et al. 2012), was first used to PCR-amplify potential 16S rRNA gene fragments from the individual metagenomes of the 25 discrete sediment samples explored along SSK42/5 and SSK42/6. For a given sediment sample, its duplicate metagenomic DNA preparations were mixed in equal proportions before performing the PCR. The 25 different PCR-products obtained in this way were all approximately 560 bp long, so were deemed to be unfit for sequencing by the 400 nucleotide read chemistry of the Ion S5 platform. Accordingly, all the PCR products were used as templates to perform nested PCR using the *Archaea-*specific primer pair A571F (5’-GCYTAAAGSRICCGTAGC-3’) and Ab909R (5’-TTTCAGYCTTGCGRCCGTAC-3’) that amplify the V4-V5 overlapping region of all archaeal 16S rRNA genes (Baker et al. 2003). This also helped eliminate all potential unspecific amplicons before the PCR-products were individually sequenced on an Ion S5 next generation DNA sequencer (Thermo Fisher Scientific, USA), using the multiplexed fusion primer protocol (Ghosh et al. 2015; Mondal et al. 2022). The sequence datasets obtained in this way were deposited in the Sequence Read Archive of National Center for Biotechnology Information (NCBI), USA, under the BioProject PRJNA309469, with BioSample and Run accession numbers given in Tables S5 and S6. Reads contained in a given 16S rRNA gene sequence dataset were trimmed for their adapters, those having lengths < 150 bp and Phred score < 20 were filtered out, and then the remaining reads were annotated taxonomically using the online utility called RDP Classifier (Wang et al. 2007).

### Detection of nitric oxide dismutase genes within the assembled metagenomes of SSK42/5 and SSK42/6

Microbial dismutation of nitric oxide (NO) can be a potential source of O_2_ within dark ecosystems (Ettwig et al. 2010; Kraft et al. 2022; Ruff et al. 2023). The NO dismutase (Nod) enzyme of bacteria such as *Candidatus* Methylomirabilis oxyfera, and strain HdN1, reportedly convert NO directly to N_2_ and O_2_ (Ettwig et al. 2010) whereas its homologs putatively present in ammonia (NH_3_)-oxidizing archaea such as *N*. *maritimus* apparently carry out the same conversion via nitrous oxide (N_2_O) formation (Kraft et al. 2022). To check whether there was any feasibility of NO-derived O_2_ sustaining the aerobic bacteria of SSK42/5 and SSK42/6, every contig of each assembled metagenome obtained along the two sediment-cores was used as a query sequence to search for genes encoding nitric oxide dismutase, using the blastx algorithm (Camacho et al. 2009). For this purpose translated amino acid sequences of *nod* gene homologs from *Candidatus* Methylomirabilis oxyfera (NCBI accession numbers ANF28199.1, APP93282.1, ANF28196.1 and APP93283.1) were used as the subject database (notably, in public databases, no Nod sequence is available thus far from ammonia-oxidizing archaea). BLAST+ executables, a suite of command-line programs to run Basic Local Alignment Search Tool, available at the website of NCBI, was used for the aforesaid searches, and hits having more than 45 nucleotide alignment, less than 10^-6^ e value, and more than 40 percent identity were reported.

## RESULTS

### Aerobic chemoorganotrophs from ASOMZ sediments

Bacterial consortia were separately enriched from the bulk (pooled) sediment samples of the two cores SSK42/5 and SSK42/6 via aerobic incubation in three distinct chemoorganoheterotrophic media, leading to the isolation of an overall 39 pure culture strains: 17 from SSK42/5 and 22 from SSK42/6 (Table 1) that appeared to be distinct in terms of colony morphology and texture. 30 of these were obtained on the copiotrophic ASW_HY medium that contained 0.5 g L^-1^ yeast extract as a source of complex organic carbon, 5 strains were obtained on the oligotrophic ASW_LY medium that contained low concentration (0.001 g L^-1^) of yeast extract, and 4 were retrieved on the synthetic BBB_A medium that had acetate (10 mM) as the sole organic carbon source. 16S rRNA gene sequence-based taxonomic classification clustered the 39 isolates into nine species-level groups, which in turn were ascribable to eight genera: *Brevibacterium*, *Brucella*, *Gordonia*, *Halomonas*, *Halopseudomonas*, *Marinobacter*, *Mesobacillus* and *Stenotrophomonas*. Two species-level groups were found to represent two separate species of *Brevibacterium*; one representative isolate from each of the nine species-level clusters was chosen for downstream experiments (Table 1). On the basis of their 16S rRNA gene sequence similarities (Table 1), conserved bacterial marker gene sequence phylogeny (Fig. 1), and overall genome-genome relatedness (dDDH values, Table 1), with closely related bacterial species, the representative isolates BDJS002 and JSBI002 were identified as two distinct and potentially novel species of the genus *Brevibacterium*. In the same way, the representative isolates JSBI001, SMJS1, BDJS001, SMJS2, AN1, SBJS01 and SBJS02 were classified as *Brucella* sp., *Gordonia hongkongensis*, *Halomonas* sp., *Halopseudomonas bauzanensis*, *Marinobacter segnicrescens*, *Mesobacillus* sp. and *Stenotrophomonas* sp., respectively (Fig. 1, Table 1).

**Figure 1.**
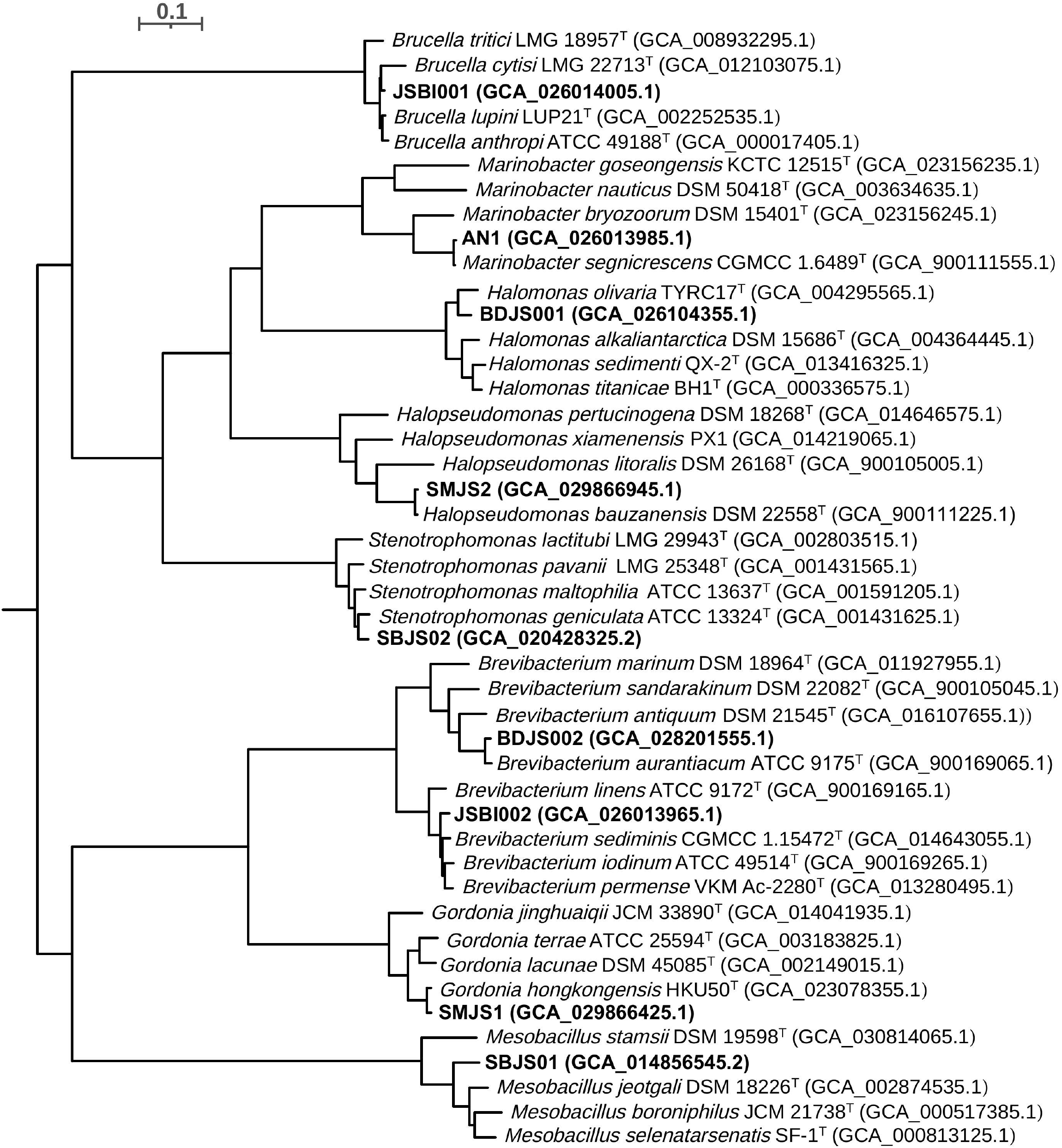
Phylogenetic tree (with 10000 bootstrap tests) constructed based on 92 universal bacterial core genes, showing the relationships among the ASOMZ-sediment strains and their closest relatives identified based on rRNA gene sequence similarities and/or dDDH. The scale bar shown indicates 10% nucleotide substitution.

**Table 1.**
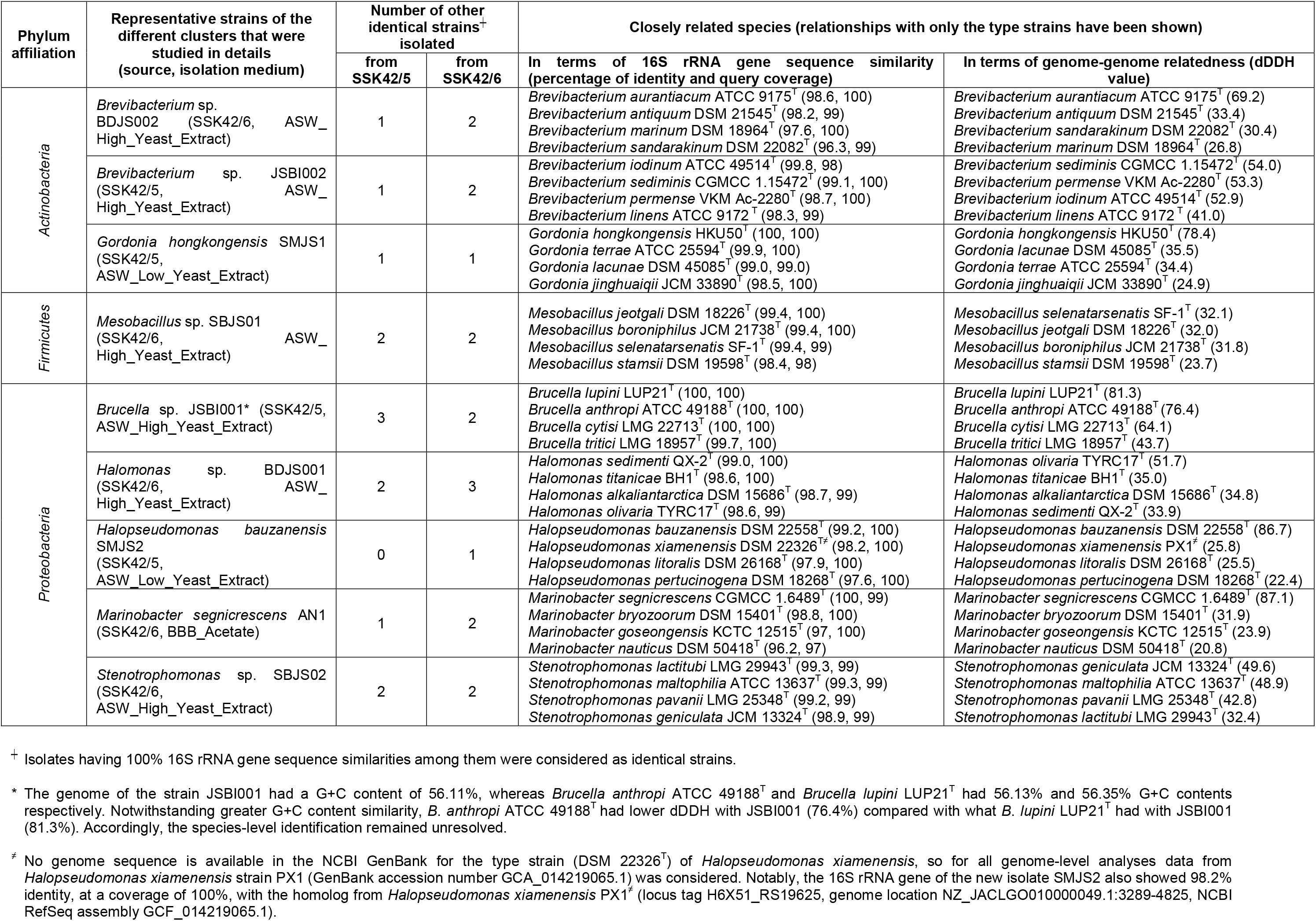
Isolation and identification of the chemoorganoheterotrophic bacterial strains isolated from Arabian Sea OMZ sediments.

### Obligately aerobic nature of all but two new isolates

All the nine representative strains could grow aerobically [at 21% (v/v) O_2_] in ASW_HY medium, with their CFU counts increasing to 0.6-3.9 × 10^4^ percent of the corresponding initial levels, after 1-2 days of incubation (Fig. 2A, Table S7). They were therefore maintained routinely in ASW_HY under aerobic condition. All the new isolates could also grow aerobically in ASW supplemented with 10 mM sodium acetate (ASW_A), with CFU counts increasing to 0.5-1.7 × 10^4^ percent of the corresponding initial levels, after 1-2 days of incubation (Table S7).

**Figure 2.**
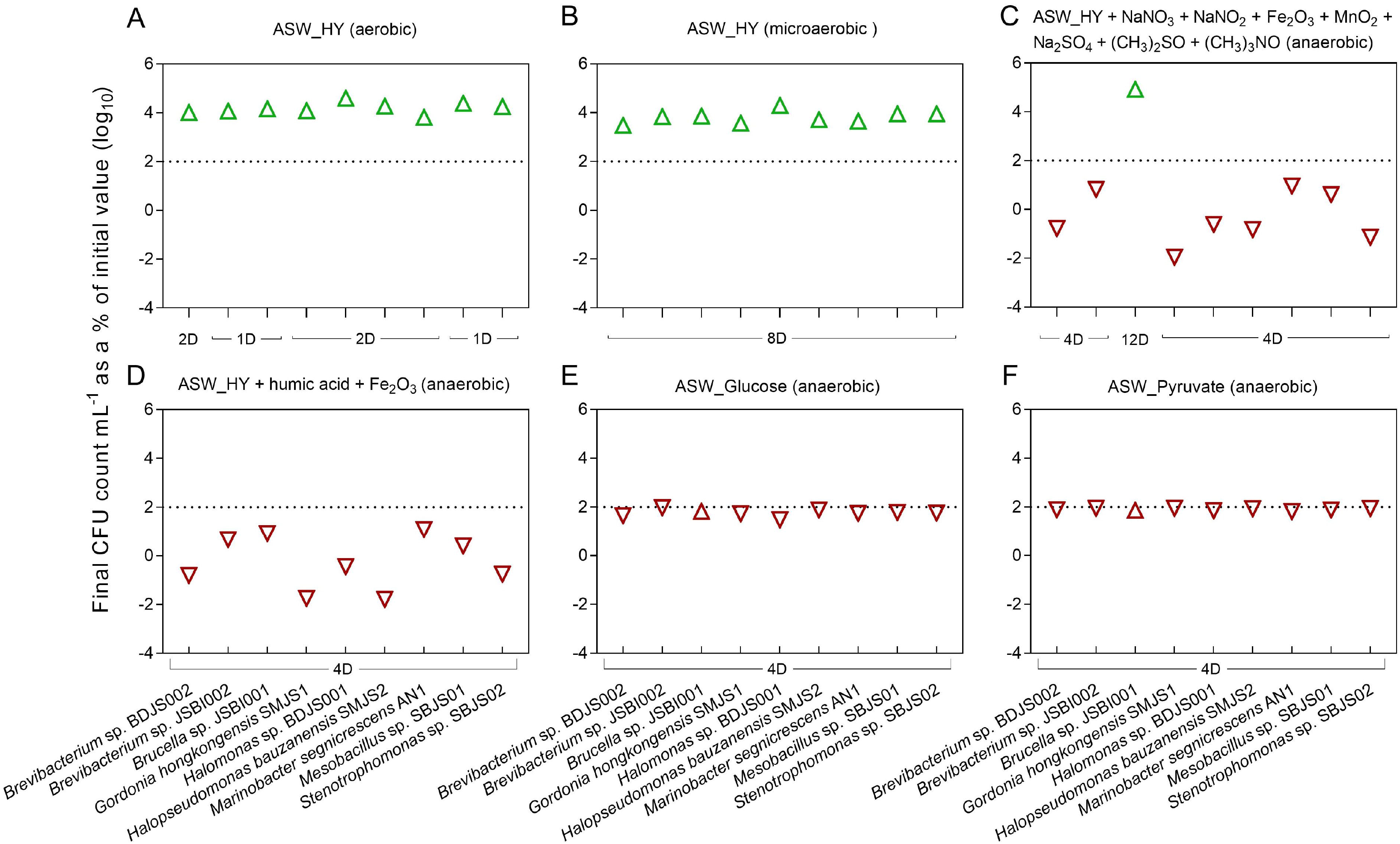
Growth (indicated by green triangles) or survival (indicated by red triangles) of the nine OMZ-sediment isolates in different ASW-based media under aerobic, microaerobic, or anaerobic condition. (**A**) Results of incubation in ASW_HY (ASW plus high concentration of yeast extract) for one (1D) or two (2D) days under aerobic condition. (**B**) Results of incubation in ASW_HY medium for eight days (8D) days under microaerobic condition. (**C**) Results of incubation in ASW_HY supplemented with six different terminal electron acceptors for four (4D) or twelve (12D) days under anaerobic condition. (**D**) Results of incubation in ASW_HY supplemented with humic acids and Fe_2_O_3_ as the terminal electron acceptors for four days (4D) under anaerobic condition. (**E**) Results of incubation in ASW_Glucose for four days (4D) under anaerobic condition. (**F**) Results of incubation in ASW_Pyruvate for four days (4D) under anaerobic condition. Final CFU counts recorded after the individual incubation periods have been plotted along the *y*-axes of the graphs, as the percentages of the corresponding initial CFU counts of the cultures (all initial and final CFU values underlying the six scatter plots are given in Table S7). All the percentage values plotted are arithmetic means of equivalent data obtained from three experimental replicates; standard deviation was < 2% of the mean for all the datasets in question. For all the percentage values in question, their Log_10_ transformations have been plotted to resolve the wide range of the values within a single frame. In each graph, the dotted grid line at the level of two 2 indicates the initial level (100%) of CFU count of the cultures concerned.

None of the ASOMZ-sediment isolates except *Brucella* sp. JSBI001 could grow anaerobically [at zero partial pressure of O_2_] in ASW_HY supplemented with (CH_3_)_3_NO, (CH_3_)_2_SO, Fe_2_O_3_, MnO_2_, NaNO_3_, NaNO_2_ and Na_2_SO_4_. Cultures of the eight test strains belonging to the genera *Brevibacterium*, *Gordonia*, *Halomonas*, *Halopseudomonas*, *Marinobacter*, *Mesobacillus* and *Stenotrophomonas*, upon anaerobic incubation in ASW_HY supplemented with all the seven electron acceptors together, retained only 0.01-9.2 percent in their initial CFU counts after 4 days (Fig. 2C); no CFU was left in any of the eight cultures after 8 days (Table S7). On the other hand, after 12 days of incubation under these conditions, CFU count of *Brucella* sp. JSBI001 increased to 8.3 × 10^4^ percent of the initial (Fig. 2C, Table S7). After anaerobic incubation for 12 days in ASW_HY supplemented with NaNO_3_, or NaNO_2_, CFU count of JSBI001 increased to 1.1 × 10^4^ percent, and 9.3 × 10^3^ percent, of the initial values respectively (Table S7). Corroboratively, 3.9 mM NaNO_3_ and 3.4 mM NaNO_2_ were found to be depleted in the two cultures, as measured by ion chromatography of the spent medium. In contrast to the above data, individual supplementation of ASW with (CH_3_)_3_NO, (CH_3_)_2_SO, Fe_2_O_3_, MnO_2_, or Na_2_SO_4_ caused the retention of no CFU in JSBI001 cultures after 8 days of anaerobic incubation (time-course data given in Table S7). Furthermore, none of the nine new isolates characterized in details, including *Brucella* sp. JSBI001, could render anaerobic growth (Fig. 2D) or maintain a fraction of the cell population in viable state (Table S7) in ASW_HY supplemented simultaneously with Fe_2_O_3_ and humic acids.

Subsequent to the above experiments, the new isolates were tested for their abilities to grow anaerobically in ASW_A medium. None of the OMZ-sediment isolates except *H*. *bauzanensis* SMJS2 could grow in the absence of O_2_ when ASW_A was collectively supplemented with the electron acceptors (CH_3_)_3_NO, (CH_3_)_2_SO, Fe_2_O_3_, MnO_2_, NaNO_3_, NaNO_2_ and Na_2_SO_4_. The eight test strains belonging to the genera *Brevibacterium*, *Brucella*, *Gordonia*, *Halomonas*, *Marinobacter*, *Mesobacillus* and *Stenotrophomonas*, when incubated in ASW_A supplemented with all the seven electron acceptors collectively, retained only 0.006-0.93 percent of their initial CFU counts after 4 days of incubation; no CFU was left in any of them after 8 days (Table S7). In contrast, the *H*. *bauzanensis* SMJS2 culture, after 12 days of incubation under these conditions, increased its CFU count to 2.1 × 10^4^ percent of the initial level (Table S7). Furthermore, SMJS2, after anaerobic incubation for 12 days in ASW_A supplemented with NaNO_2_, increased its CFU count to 2.0 × 10^4^ percent of the initial value (Table S7); corroboratively, ion chromatography showed that 3.3 mM NaNO_2_ was depleted in the SMJS2 culture. However, individual supplementation of ASW with (CH_3_)_3_NO, (CH_3_)_2_SO, Fe_2_O_3_, MnO_2_, NaNO_3_, or Na_2_SO_4_ caused the retention of no CFU in JSBI001 cultures after 8 days of anaerobic incubation (detailed data given in Table S7). Furthermore, none of the nine new strains, including *H*. *bauzanensis* SMJS2, could render anaerobic growth or survival in ASW_A supplemented simultaneously with Fe_2_O_3_ and humic acids (Table S7).

### Microaerobic growth and fermentative survival

Ability of the ASOMZ-sediment isolates to grow or survive under microaerobic condition [0.1% (v/v) O_2_] was checked in ASW_HY medium. After incubation for 4 days under such conditions within the H35 Hypoxystation, CFU count of the test strains increased 0.7-3.7 × 10^3^ percent (Table S7), while after 8 days the increase was found to be 0.3-2.0 × 10^4^ percent for the different isolates (Fig. 2B, Table S7).

The ASOMZ-sediment isolates were tested for their potentials to grow or survive amid anoxia using fermentative metabolism. After anaerobic incubation for 4 days in ASW containing glucose or pyruvate, the strains retained 30.9% to 96.2%, and 66.7% to 89.5%, of their initial CFUs respectively (Fig. 2E and 2F); after 8 days, 15.2% to 54.7%, and 9.4% to 85.6% CFUs were retained respectively (Table S7). Notably, even after 64 days of anaerobic incubation in glucose or pyruvate supplemented ASW, CFU count of the culture did not come down to zero for any of the strains in question (Table S7). To confirm the role of glucose and pyruvate (as fermentative substrates) in the above survival process, anaerobic incubation was carried for all the nine strains in ASW only (i.e. without any energy and carbon source). After 8 days of anaerobic incubation in ASW, CFU count in the culture of all but one isolate came down to zero. The anaerobic ASW culture of *Brucella* sp. JSBI001, despite retaining 1.2 × 10^3^ CFU mL^-1^ after 8 days, contained no CFU after 16 days of incubation (Table S7).

### Growth on complex carbon compounds under aerobic and microaerobic, but not anaerobic, conditions

The new isolates were tested for their abilities to grow aerobically, microaerobically, or anaerobically (Fig. 3, Tables 2 and S8), using agar, alginate, benzoate, carrageenan, cellulose, chitosan, pectin, starch or xylan as the sole carbon source in ASW-based chemoorganoheterotrophic medium.

**Figure 3.**
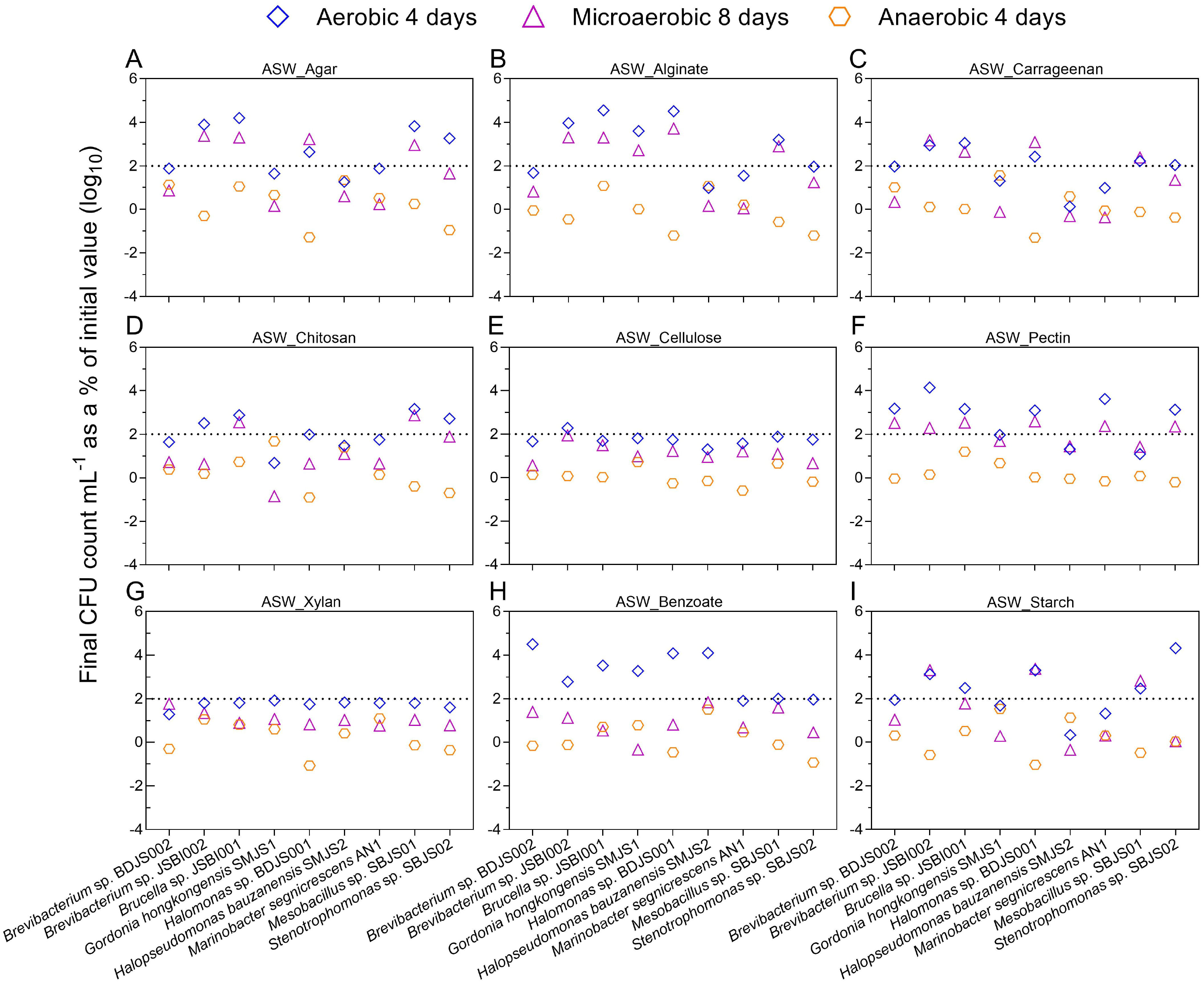
Growth or survival of the OMZ-sediment isolates in ASW-based media supplemented with different complex carbon compounds, after incubations for four, eight, and four days, under aerobic, microaerobic, and anaerobic conditions, respectively. Results of incubation in ASW supplemented with (**A**) Type-I agar, (**B**) sodium alginate, (**C**) carrageenan, (**D**) chitosan, (**E**) sodium carboxymethylcellulose, (**F**) pectin, (**G**) xylan, (**H**) sodium benzoate, and (**I**) starch have been shown in the different panels. Final CFU counts recorded after the individual incubation periods have been plotted along the *y*-axes of the graphs, as the percentages of the corresponding initial CFU counts of the cultures (all initial and final CFU values underlying the different scatter plots are given in Table S8). All percentage values shown are arithmetic means of equivalent data obtained from three experimental replicates; standard deviation was < 2% of the mean for all the datasets in question. For all percentage values, Log_10_ transformations have been plotted to resolve the wide range of the values. In each graph, the dotted grid line at the level of two 2 indicates the initial level (100%) of CFU count of the cultures.

After aerobic incubation for 4 days, benzoate and pectin were utilized for growth by six strains each; agar, alginate, carrageenan and starch were utilized by five strains each; chitosan was utilized by four strains; cellulose was utilized by only one strain; and xylan was utilized by none (Fig. 3). At the strain level, *Brevibacterium* sp. JSBI002 could utilize eight out of the nine complex carbon compounds tested for growth; *Brucella* sp. JSBI001 could utilize seven, *Halomonas* sp. BDJS001 could utilize six, while *Mesobacillus* sp. SBJS01 and *Stenotrophomonas* sp. SBJS02 could utilize 5 substrates each for growth. *Brevibacterium* sp. BDJS002 could utilize only pectin and benzoate, *G*. *hongkongensis* SMJS1 utilized only alginate and benzoate, whereas *M*. *segnicrescens* AN1 and *H*. *bauzanensis* SMJS2 could utilize only pectin and benzoate respectively (Table 2).

**Table 2.**
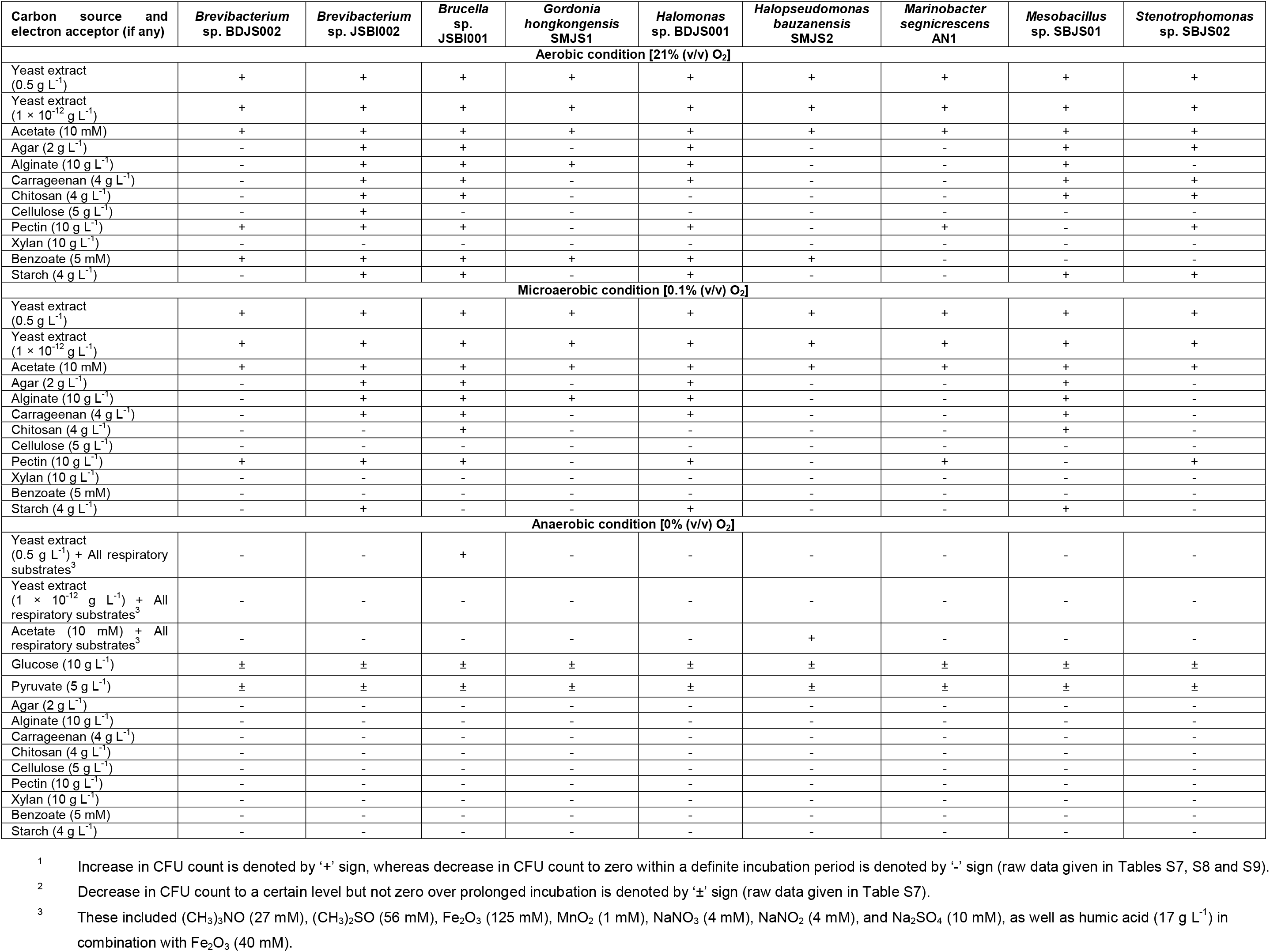
Growth^1^ or survival^2^ of the ASOMZ-sediment isolates on different organotrophic substrates under different levels of O_2_ availability.

As observed after microaerobic incubation for 8 days, pectin was utilized for growth by six strains, alginate was utilized by five strains, agar and carrageenan were utilized by four strains each, starch was utilized by three strains, while chitosan was utilized by two; cellulose, benzoate and xylan were utilized by none (Fig. 3). At the strain level, *Brevibacterium* sp. JSBI002 and *Brucella* sp. JSBI001 each could utilize five out of the nine complex carbon compounds tested, *Halomonas* sp. BDJS001 and *Mesobacillus* sp. SBJS01 could utilize three each, *M*. *segnicrescens* AN1 and *Stenotrophomonas* sp. SBJS02 utilized only pectin, *G*. *hongkongensis* SMJS1 utilized only alginate, while *Brevibacterium* sp. BDJS002 and *H*. *bauzanensis* SMJS2 could utilize none (Table 2).

After anaerobic incubation for 8 days, none of the test strains could use agar, alginate, benzoate, carrageenan, cellulose, chitosan, pectin, starch or xylan as the sole carbon source to grow chemoorganoheterotrophically in ASW containing (CH_3_)_3_NO, (CH_3_)_2_SO, Fe_2_O_3_, MnO_2_, NaNO_3_, NaNO_2_ and Na_2_SO_4_; or Fe_2_O_3_ and humic acids together (Fig. 3, Tables 2 and S8).

### Cell density dependent extreme oligotrophy under aerobic and microaerobic, but not anaerobic, conditions

The ASOMZ-sediment isolates were tested for their abilities to grow aerobically, microaerobically, or anaerobically, in ASW containing 1 × 10^-12^ g L^-1^ yeast extract. Irrespective of the oxygen concentration provided, no growth was observed in any of the nine strains characterized in this study when the initial cell count of the culture was set at ∼10^6^ CFU mL^-1^ (Fig. 4A, Tables 2 and S9). In contrast, starting with an initial number of ∼10^3^ CFU mL^-1^, the CFU counts of the different strains increased to 0.06-1.3 × 10^4^ percent or 0.5-6.7 × 10^4^ percent of the corresponding initial levels, after 8 days of incubation, under aerobic or microaerobic condition respectively; subsequently, the CFU count remained static even after incubation for several weeks (Fig. 4B, Tables 2 and S9). Low initial cell density, however, could not facilitate oligotrophic growth under anaerobic condition (Fig. 4B, Tables 2 and S9).

**Figure 4.**
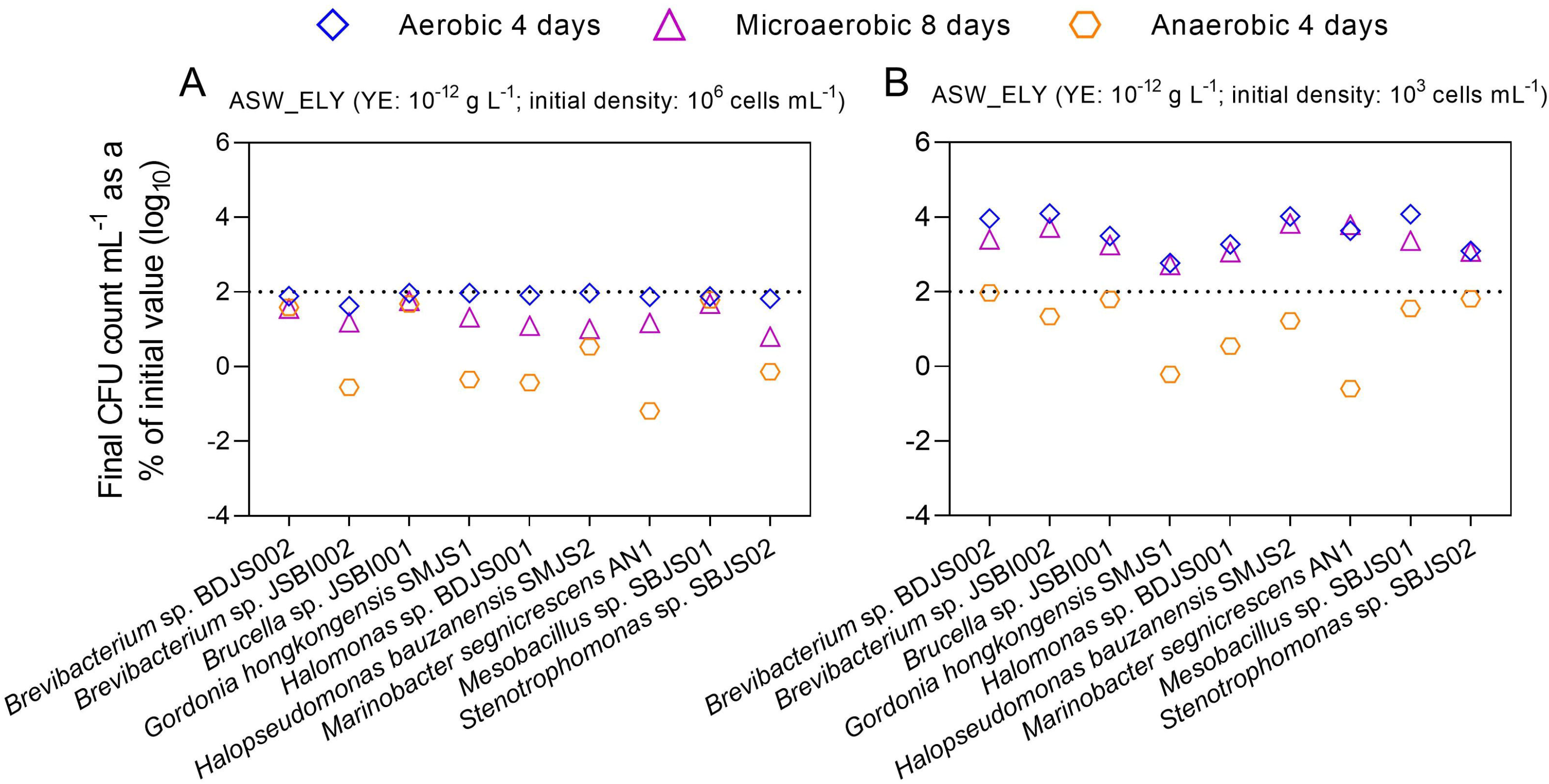
Growth or survival of the OMZ-sediment isolates in ASW-based media supplemented with extremely low concentration of yeast extract (ASW_ELY) after incubations for four, eight, and four days, under aerobic, microaerobic, and anaerobic conditions, respectively. Results of incubation with (**A**) relatively higher (∼10^6^ CFU mL^-1^ of culture) and (**B**) lower (∼10^3^ CFU mL^-1^ of culture) initial cell density have been shown in the two panels. Final CFU counts recorded after the individual incubation periods have been plotted along the *y*-axes of the graphs, as the percentages of the corresponding initial CFU counts of the cultures (all initial and final CFU values underlying the two scatter plots are given in Table S9). All percentage values shown are arithmetic means of equivalent data obtained from three experimental replicates; standard deviation was < 2% of the mean for all the datasets in question. For all percentage values, Log_10_ transformations have been plotted to resolve the wide range of the values. In each graph, the dotted grid line at the level of two 2 indicates the initial level (100%) of CFU count of the cultures.

### A brief overview of the genome contents of the ASOMZ-sediment isolates

*De novo* hybrid assembly of short and long reads, generated from Ion S5 and MinION sequencing platforms respectively, yielded a single closed circular chromosome for each of the nine new isolates (Table S2). In addition to these, *Brucella* sp. JSBI001 and *H*. *bauzanensis* SMJS2 were found to have one closed circular plasmid each, while *G*. *hongkongensis* SMJS1 had two such replicons (Table S2). The genomes of the new isolates had considerable variations in terms of their gene contents as well as their patterns of CDS distribution across the various functional categories of clusters of orthologous genes (COGs; Fig. S1; Table S2). However, the chromosomes of *M*. *segnicrescens* AN1, *H*. *bauzanensis* SMJS2, *Stenotrophomonas* sp. SBJS02 and *Mesobacillus* sp. SBJS01 on one hand, and those of *Halomonas* sp. BDJS001, *Brucella* sp. JSBI001, *Brevibacterium* sp. JSBI002, *Brevibacterium* sp. BDJS002 and *G*. *hongkongensis* SMJS1 on the other, had mutually similar patterns of CDS distribution across the various COG categories (heat map shown in Fig. S2). Likewise, the two plasmids of *G*. *hongkongensis* SMJS1, together with the plasmid of *H. bauzanensis* SMJS2, appeared to be mutually similar, and at the same time distinct from the plasmid of *Brucella* sp. JSBI001.

### Genes conferring bioenergetic versatility to the ASOMZ-sediment strains

Genome analysis revealed the presence of low as well as high O_2_-affinity cytochrome oxidases in all the nine strains characterized from ASOMZ sediments. Genes for *aa3*-type cytochrome oxidase and cytochrome-*bd* ubiquinol oxidase were detected in all the nine strains, while those for *cbb3*-type cytochrome oxidase were identified in *Brucella* sp. JSBI001, *Halomonas* sp. BDJS001, *H*. *bauzanensis* SMJS2 and *M. segnicrescens* AN1 (Fig. 5, Table S10).

**Figure 5.**
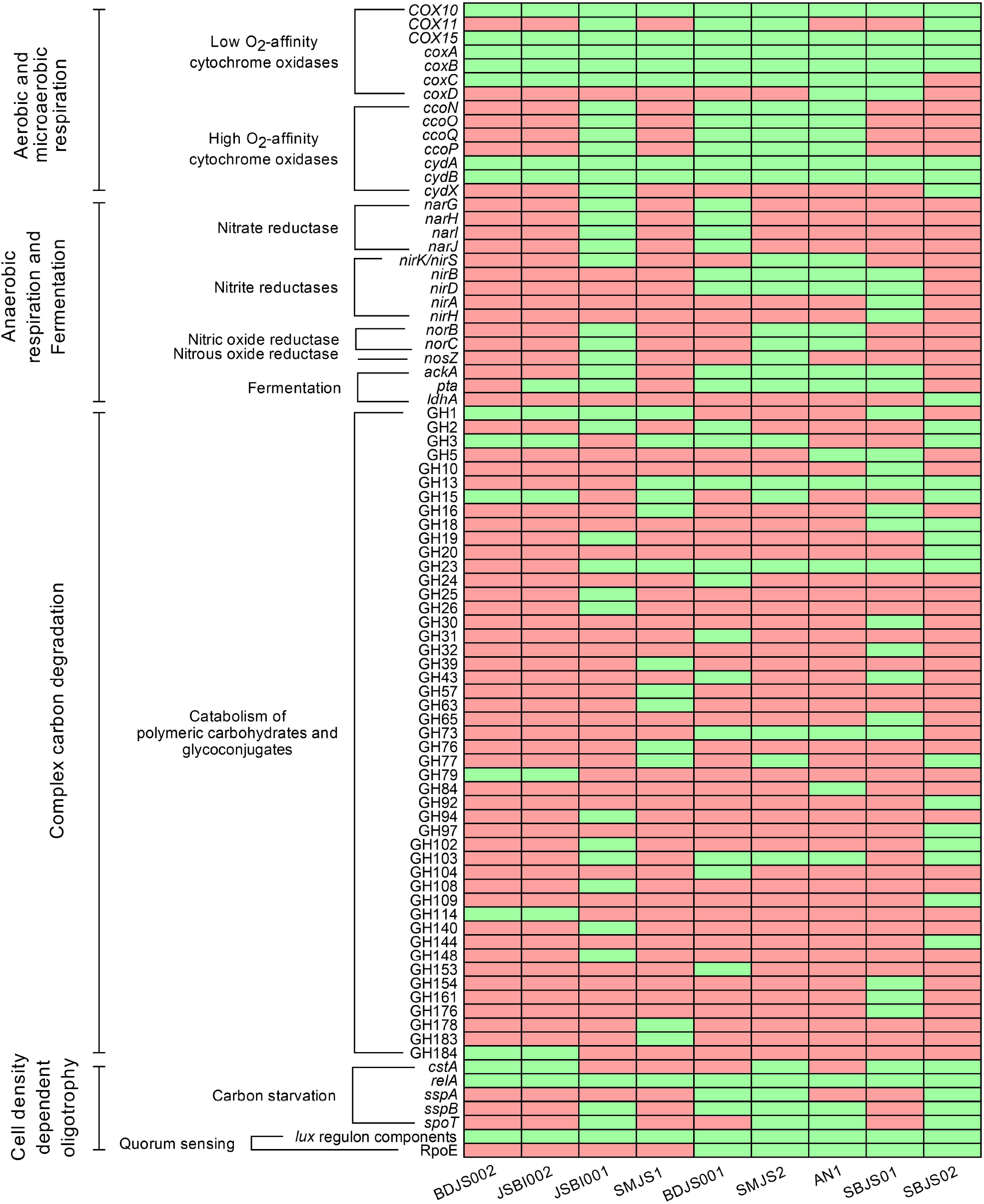
Presence (indicated by green boxes) or absence (indicated by red boxes) of genes (CDSs) for some of the key enzymes of aerobic and microaerobic respiration, anaerobic respiration and fermentation, catabolism of polymeric carbohydrates and glycoconjugates, and cell density dependent oligotrophy, within the annotated genomes of the ASOMZ-sediment isolates.

*Brucella* sp. JSBI001, the only isolate capable of anaerobic growth via nitrate and nitrite respiration (Table S7), was found to contain all genes that are required for complete denitrification, i.e. reduction of nitrate to dinitrogen (Fig. 5, Table S11). The *H*. *bauzanensis* strain SMJS2, which could perform anaerobic growth via nitrite respiration alone (Table S7), contained all the genes needed for the dissimilatory reduction of nitrite to dinitrogen or ammonia (Fig. 5, Table S11).

Although none of the other ASOMZ-sediment isolates exhibited nitrate or nitrite respiration in ASW_HY and ASW_A (Table S7), three of them possessed genes for the dissimilatory reduction of nitrate and/or nitrite. The genome of *Halomonas* sp. BDJS01 contained all the oxidoreductase genes that are required to reduce NO_3_ to NH_3_; the *M*. *segnicrescens* AN1 genome encoded all the enzymes required for the reduction of NO to N_2_O or NH_3_, while *Mesobacillus* sp. SBJS01 encoded two distinct enzyme systems for NO reduction to NH_3_ (Fig. 5, Table S11).

Genes encoding phosphate acetyltransferase, acetate kinase, and lactate dehydrogenase were detected in the genomes of all the new isolates except *Brevibacterium* sp. BDJS002, *G*. *hongkongensis* SMJS1 and *Stenotrophomonas* sp. SBJS02 (Fig. 5, Table S11); all these enzymes are known to be central to fermentative metabolism and energy generation under anaerobic condition.

### Genes for complex organic matter degradation in the new isolates

All the new isolates were found to possess an array of genes to encode enzymes that can break the different glycosidic bonds of diverse polymeric carbohydrates and glycoconjugates. Across the nine strains, a sum total of 145 such genes were detected which coded for carbohydrate-active enzymes classifiable under 47 distinct glycoside hydrolase (GH) families (Fig. 5, Table S12). Member proteins of the GH families detected are known to render - among other functions - β-glucosidase, β-galactosidase, endo-α-1,4-galactosaminidase, endo-β-1,3-xylanase, endo-β-1,4-glucanase, α-amylase, oligo-α-1,6-glucosidase, and chitinase activities, which in turn are central to the catabolism of the different homo- and hetero-polysaccharides on which the new isolates were found to grow chemoorganoheterotrophically (Fig. 3, Table 2). So far as the individual ASOMZ-sediment strains are concerned, *Stenotrophomonas* sp. SBJS02 possessed the highest (26) number of glycoside hydrolase genes distributed over the most number (16) of GH families. On the other hand, the two *Brevibacterium* strains BDJS002 and JSBI002 had the fewest (7) glycoside hydrolase genes distributed over the lowest number (6) of GH families (Fig. 5, Table S12).

So far as the degradation of aromatic compounds is concerned, only *Halomonas* sp. BDJS001 and *H*. *bauzanensis* SMJS2 were found to have all the genes necessary for the catabolism of benzoate to catechol, whereas *Halomonas* sp. BDJS001 alone possessed all the genes needed for catechol ortho-cleavage; *Brevibacterium* sp. BDJS002 and *H*. *bauzanensis* SMJS2 possessed only some of the genes that are required for catechol ortho-cleavage (Fig. S3, Table S13). Furthermore, all the isolates encompassed one or more, but not all, genes for catechol meta-cleavage. Whereas only *Brevibacterium* sp. BDJS002 contained the entire repertoire genes needed for cyclohexanecarboxylic acid catabolism, others had none (Fig. S3, Table S13).

### Genomic bases of cell density dependent oligotrophy in the new isolates

Genes coding for proteins concerned with signal transduction in response to carbon starvation were found to be present in all the nine strains characterized (Fig. 5, Table S14). Furthermore, all the nine genomes encompassed one or more genes to code for protein mediators of stringent response to nutrient-level fluctuation. All the nine genomes analyzed encompassed a large number of genes for quorum sensing, including those attributed to the Lux regulon (Fig. 5, Table S14). In addition, the genomes of *Halomonas* sp. BDJS001, *H*. *bauzanensis* SMJS2, *M. segnicrescens* AN1, *Mesobacillus* sp. SBJS01 and *Stenotrophomonas* sp. SBJS02 were found to encode the alternative sigma factor RpoE, which mediates cellular response to diverse environmental stressors including starvation.

### The chemoorganotrophic strains are ubiquitous and active across the sediment horizon explored

When all quality-filtered metagenomic reads available for each of the 25 discrete sediment samples of SSK42/5 and SSK42/6 (Table S3) were mapped individually onto the genomes of the nine strains one by one, sizeable percentages of reads from the different metagenomic datasets mapped with considerable coverage breadths and depths onto each of the target genomes (Table S15). As an exception, only for the metagenome of the 190 cmbsf sample of SSK42/5 no read mapped onto the *Brevibacterium* strains BDJS002 and JSBI002. In SSK42/5, cumulative abundance of these organisms remained largely consistent (0.1% to 0.3% of a metagenome) from the core-top till the core-bottom, except for the two spikes at 45 cmbsf and 90 cmbsf where these strains accounted for 1.3% and 1.1% of the metagenome (Table S15). In SSK42/6, however, overall representation of these organisms in the metagenome decreased from the core-top until 135 cmbsf (from 2.3% to 0.5% of the metagenome), then increased (up to 11.3%) until 250 cmbsf, and eventually decreased again to 1.1% at 275 cmbsf sediment-depth (Table S15).

With regard to the individual isolates, maximum percentage of metagenomic reads mapped onto the genome of *M*. *segnicrescens* AN1, in case of almost all the sediment samples analyzed across the two cores. 0.01% to 8.2% of reads from 24 out of the 25 metagenomes of SSK42/5 and SSK42/6 mapped individually onto the AN1 genome with coverage breadths of 1.6% to 95.5% of the entire genome length of the strain, and average coverage depths of 1.2X to 12.7X (coverage depth denotes what number of metagenomic reads from a given sediment sample mapped, on an avergae, to each locus of alingmnet within a particular genome). Only for 275 cmbsf of SSK42/6, most metagenomic reads (0.8% of the total) mapped onto the genome of *Halomonas* sp. BDJS001 with a coverage breadth of 100% and average coverage depth of 2.1X (Table S15). Apart from *M*. *segnicrescens* AN1, four strains, namely *Halomonas* sp. BDJS001, *H*. *bauzanensis* SMJS2, *Stenotrophomonas* sp. SBJS02 and *Brucella* sp. JSBI001, had overall higher representation in the metagenomes, compared with the other ASOMZ-sediment isolates (Table S15).

When all quality-filtered mRNA-specific metatranscriptomic reads (Table S4) available for the surface and subsurface sediments of SSK42/5 (0 cmbsf and 15 cmbsf) and SSK42/6 (2 cmbsf and 15 cmbsf) were mapped individually onto the genomes of the new isolates one by one, substantial proportions of read from the different metatranscriptomic datasets mapped with considerable coverage breadths and depths onto each of the genomes targeted (Table S16). Collectively, the genomes of the nine strains accounted for approximately 16.6% and 9.7% of the 0 cmbsf and 15 cmbsf metatranscriptomes of SSK42/5, and 19.8% and 8.9% of the 2 cmbsf and 15 cmbsf of SSK42/6, respectively. Subsequently, to check what proportion of the metabolically active sedimentary community near a core-bottom was made up of the new isolates, reads from the metatranscriptomic dataset available for the sediment sample from 275 cmbsf of SSK42/6 were mapped onto the genomes of the new isolates. Collectively, the nine genomes accounted for approximately 20.7% of this metatranscriptome. As for the individual isolates, maximum percentage of metatranscriptomic reads mapped onto the genome of *M*. *segnicrescens* AN1, in case of all the five sediment samples analyzed (Table S16). Across the metatranscriptomes, 3.5% to 10.6% reads mapped individually onto the genome of AN1 with coverage breadths of 1.8% to 26.3% and average coverage depths of 499.7X to 1883.5X (here, coverage depth denoted what number of metatranscriptomic reads from a given sediment sample mapped, on an avergae, to each locus of alingmnet within a particular genome). Three other strains, namely *Halomonas* sp. BDJS001, *H*. *bauzanensis* SMJS2 and *Stenotrophomonas* sp. SBJS02, had overall higher representations in the metatranscriptomes, compared with the other ASOMZ-sediment isolates (Table S16).

### *In situ* expression of the metabolisms that characterized the ASOMZ-sediment isolates

The metatranscriptomic datasets available for the two surface-sediment samples (0 cmbsf and 2 cmbsf for SSK42/5 and SSK42/6 respectively), two subsurface sediment samples (15 cmbsf for both the core), and one deep-sediment sample (275 cmbsf from SSK42/6), were individually assembled *de novo*; subsequently, the contigs obtained in each assembly were annotated for putative CDSs (Table S4). The five gene/CDS-catalogs obtained in this way were searched for homologs of the genes that are known to govern those metabolic properties which were identified above as central to the ecological fitness of the chemoorganotrophic isolates (Fig. 6, Table S17).

**Figure 6.**
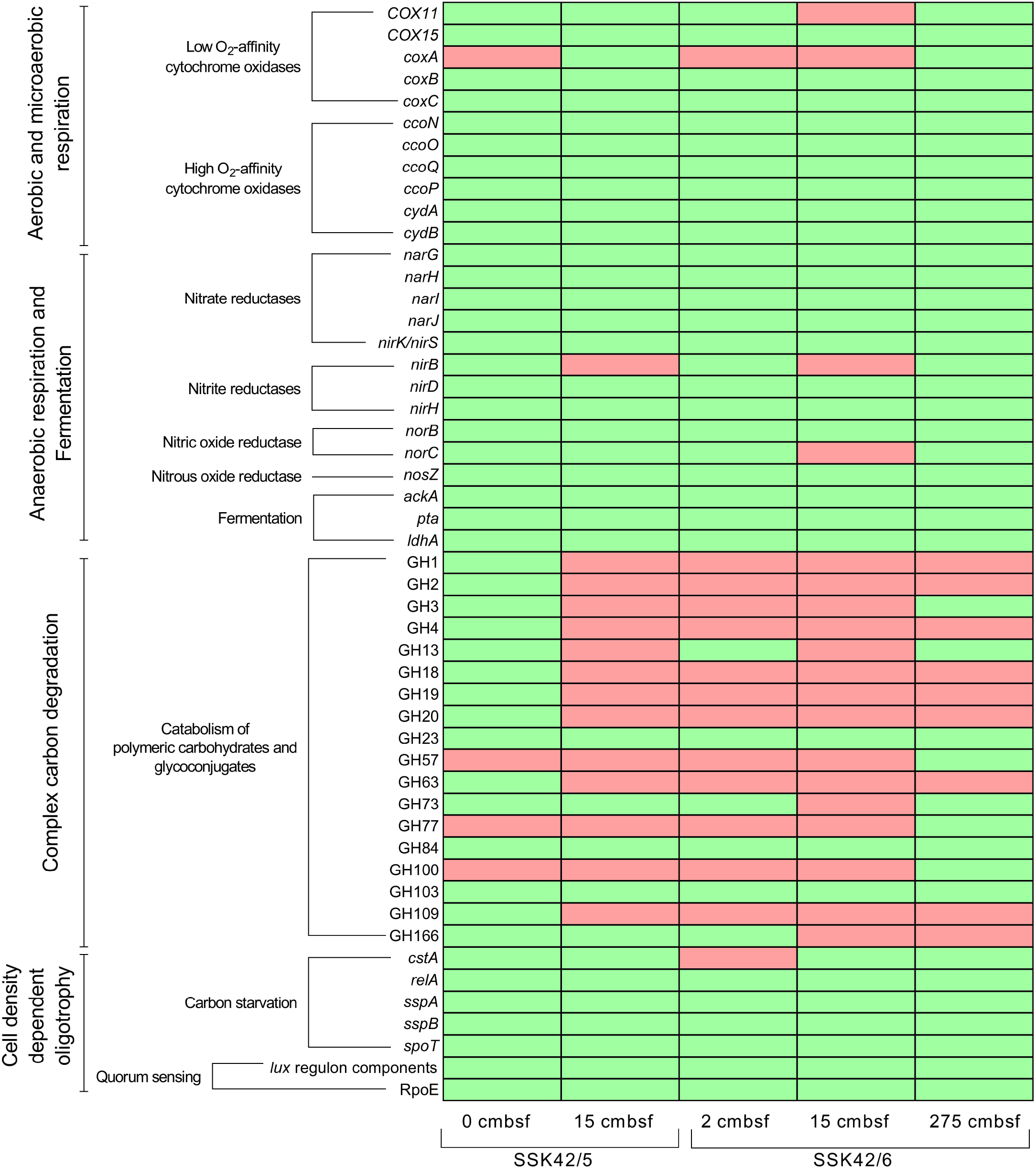
Presence (indicated by green boxes) or absence (indicated by red boxes) of genes (CDSs) for some of the key enzymes of aerobic and microaerobic respiration, anaerobic respiration and fermentation, catabolism of polymeric carbohydrates and glycoconjugates, and cell density dependent oligotrophy, within the metatranscriptomes assembled from the different sediment samples of SSK42/5 and SSK42/6.

Several genes for respiration under fully aerobic to acutely hypoxic conditions were present in all the assembled metatranscriptomes (these included genes for the high O_2_-affinity *cbb3*-type cytochrome oxidase and cytochrome-*bd* ubiquinol oxidase) alongside those for respiration via reduction of nitrate/nitrite. Genes for fermentative energy transduction (ATP generation from carbon substrates in the absence of O_2_), oligotrophy (starvation endurance), and quorum sensing (including components of the Lux regulon) were also detected in considerable numbers within all the five metatranscriptomes (Fig. 6, Table S17).

Different genes governing the catabolism of polymeric carbohydrates and glycoconjugates were identified in all the five metatranscriptomes assembled (Fig. 6, Table S17). A sum total of 141 such genes were detected across the metatranscriptomes which coded for carbohydrate-active enzymes classifiable under 18 distinct GH families. Whereas highest (51) number of glycoside hydrolase genes distributed over the most number (16) of GH families were detected in the metatranscriptome of the 0 cmbsf sediment sample from SSK42/5, lowest (3) number of glycoside hydrolase genes (these were distributed over 3 GH families) were found in the metatranscriptome of the 15 cmbsf sample from SSK42/6 (Fig. 6, Table S17).

So far as aromatic compounds degradation is concerned, genes for the transformation of benzoate to catechol were found to be present alongside those for catechol ortho-cleavage in the four surficial or subsurficial metatranscriptomes analyzed, i.e. 0 cmbsf and 15 cmbsf of SSK42/5, and 2 cmbsf and 15 cmbsf of SSK42/6. However, genes for catechol meta-cleavage, and cyclohexanecarboxylic acid utilization, were present only in the metatranscriptomes of 0 cmbsf and 15 cmbsf of SSK42/5 respectively (Fig. S4, Table S17).

### Molecular signatures of *Nitrosopumilus* and nitric oxide dismutase in the metagenomes of SSK42/5 and SSK42/6

From the physiological characterizations carried out thus far, seven out of the nine ASOMZ-sediment strains appeared to be obligately aerobic. Therefore, some explanation was required to be put forth (for subsequent testing) with regard to how those bacteria having neither anaerobic respiration capabilities nor fermentative growth potentials could be metabolically active, and relatively abundant, in these extremely reduced, sulfide-containing, sediments, which also have less probability of O_2_ influx, compared with other shelf/slope sediment systems that are situated outside of OMZ territories. Ammonia-oxidizing *Nitrosopumilus* species being known sources of biogenic O_2_ in marine environments (Walker et al. 2010; Kraft et al. 2022), their molecular footprint were searched in the context of the *in situ* survival of obligate aerobes within the microbiomes of SSK42/5 and SSK42/6.

25 distinct gene-catalogs were obtained by individually annotating the 25 assembled metagenomes of the equal number of sediment samples explored along SSK42/5 and SSK42/6 (Table S3). When these catalogs were searched for CDSs having maximum sequence similarities with homologs from species of *Nitrosopumilus*, 0.01-0.19% of the CDSs annotated for the 12 discrete samples of SSK42/5 were found to be ascribable to *N*. *maritimus* and 0.01-0.22% to *Nitrosopumilus sediminis* (Fig. 7, Table S18). Likewise, 0-0.09% of the CDSs annotated for the 13 sediment samples of SSK42/6 were ascribable to *N*. *maritimus* (the only sediment site to have no *N*. *maritimus* affiliated CDS was 250 cmbsf of SSK42/6) and 0.01-0.11% to *N*. *sediminis* (Fig. 7, Table S19).

**Figure 7.**
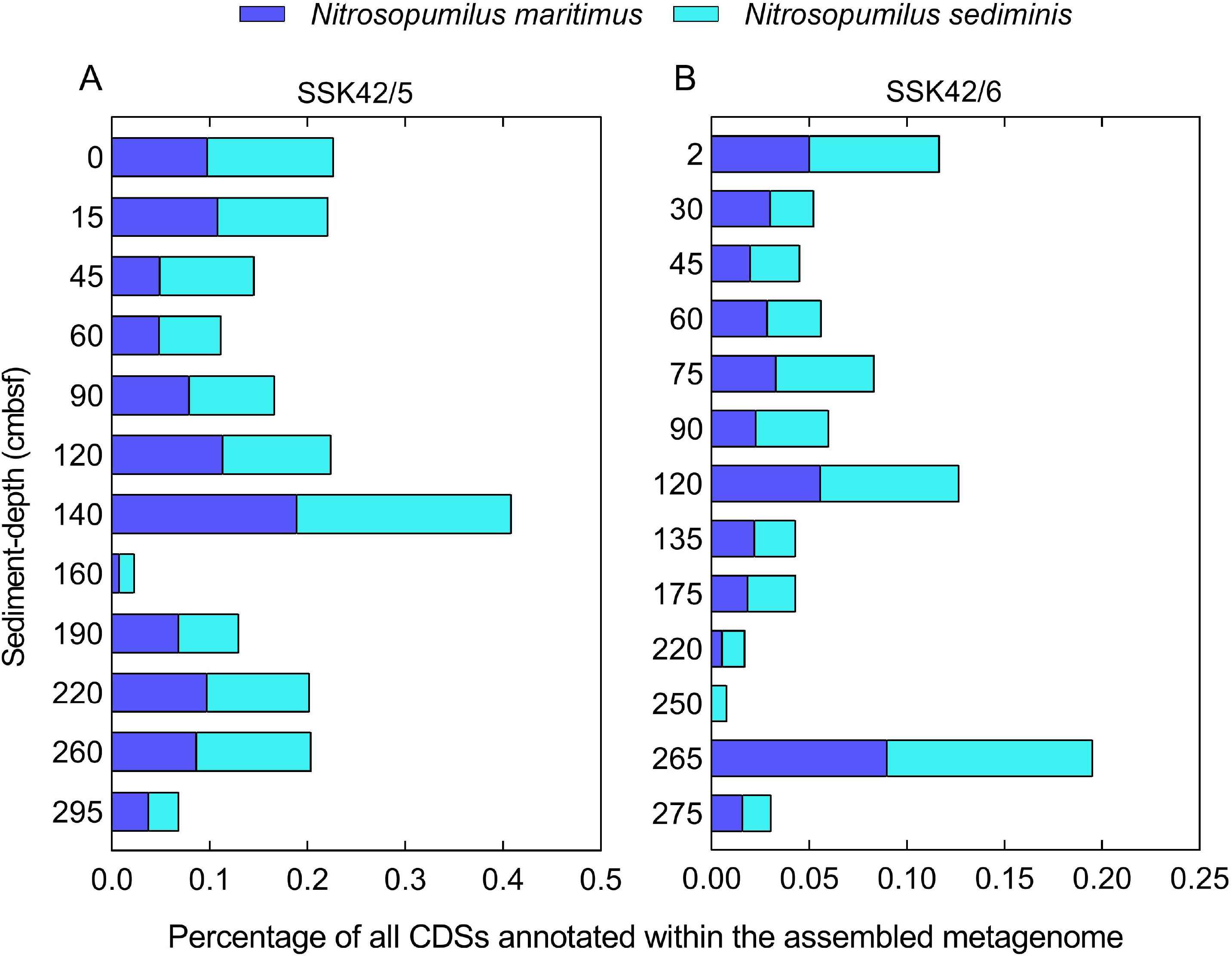
Histograms showing what percentages of the CDS contents of the different metagenomes assembled along (**A**) SSK42/5 and (**B**) SSK42/6 were ascribable to *Nitrosopumilus maritimus* and *Nitrosopumilus sediminis* on the basis of sequence similarity with homologous genes. The scales for both sediment-depth (plotted along the *y*-axes) and CDS percentage (plotted along the *x*-axes) are linear.

Assembled metagenomes of all the sediment samples of SSK42/5 and SSK42/6, except 220 cmbsf and 265 cmbsf of the latter core, were found to encompass genes encoding nitric oxide dismutase (Nod). In SSK42/5, most numbers (19 and 17) of *nod* gene homologs (CDSs) were detected at 160 cmbsf and 190 cmbsf, whereas in SSK42/6, most *nod* homologs (17 and 14) were detected at 2 cmbsf and 120 cmbsf (Table S20).

Furthermore, the metagenomic DNA preparations available for the 25 sediment samples explored across SSK42/5 and SSK42/6 were used individually as templates to PCR-amplify V4-V5 overlapping regions of 16S rRNA gene fragments using *Nitrosopumilus-*specific, followed by *Archaea-*specific, primers. When these PCR-products were sequenced and annotated, reads attributable to the genus *Nitrosopumilus* were identified in all the 25 experimental sets (Table S21).

## DISCUSSION

### Constraints faced by aerobic chemoorganoheterotrophs in the ASOMZ sediments

High levels of correspondence witnessed between the different metagenomes and metatranscriptomes of SSK42/5 and SSK42/6 on one hand and the genomes of the new isolates on the other illustrated the ubiquitous presence and activity of the strains across the ASOMZ sediment horizon explored (Tables S15 and S16). At the same time, literature survey showed that close phylogenetic relatives of the OMZ-sediment isolates were widespread in diverse marine habitats. However, the sustenance and biogeochemical activity of aerobic chemoorganoheterotrophs within the present study sites are constrained by two critical factors.

Whereas the deep subsurfaces of open ocean sites such as those studied extensively in the East (D‘Hondt et al. 2004; Batzke et al. 2007) and South (D’Hondt et al. 2009, 2015; Morono et al. 2020) Pacific Oceans remain oxic due to their characteristic geologies, sediment systems within shelf/slope territories, especially those in the upwelling regions and OMZs (including the present study sites), turn anoxic within a few meters, or even centimeters, from the seafloor, as diffusion from overlying water-columns constitute the only means of oxidant input for the latter category of marine subsurface ecosystems (Bhattacharya et al. 2019; Fernandes et al. 2022). Open ocean, as well as shelf/slope, sediments, however, are characteristically unified by the progressive limitation of bioavailable organic carbon down the sediment-depths (Arndt et al. 2013; Lever et al. 2015; Fernandes et al. 2018, 2022; Sarkar et al. 2022).

In the open ocean sites, oxidant requirements of the facultatively aerobic bacteria thriving even in the deepest sediment layers explored (these include horizons hundreds of meters below the seafloor) are sufficed by the small but definite amounts of NO, Mn^4+^, Fe^3+^ and O that are present in the ecosystems (D‘Hondt et al. 2003; Aiello et al. 2006; Batzke et al. 2007). Their extremely low metabolic (especially respiratory) rates are apparently instrumental in sustaining them for even millions of years under acute organic carbon scarcity (Jørgensen 2011; Langerhuus et al. 2012; Lomstein et al. 2012; D’Hondt et al. 2015; Braun et al. 2017). By contrast, *in situ* metabolisms of the aerobic bacterial chemoorganoheterotrophs isolated in the present study from the few thousands of years old (only a few meters below the seafloor), highly-sulfidic sediments of Arabian Sea OMZ (Fernandes et al. 2018; Bhattacharya et al. 2021) are constrained primarily by the lack of oxidants in the habitat (Breuer et al. 2009; Cavan et al. 2017; Jessen et al. 2017), even as organic carbon and energy sources also start depleting near the core-bottoms, thereby ushering an additional constraint on their survival and activity *in situ*.

### Quorum-sensing-based extreme oligotrophy as a key adaptation to carbon scarcity under aerobic and microaerobic, but not anaerobic, conditions

Although the seafloor in the ASOMZ sites SSK42/5 and SSK42/6 receives copious organic matter from the highly productive water-column, the microbiome, down the sediment-depths, gets increasingly constrained for utilizable sources of carbon and energy, as the total organic carbon content of these sediments decreases from ∼4 % (w/w) near the core-tops to 0.5-1.0 % (w/w) near the core-bottoms (Fernandes et al. 2018). Further depletion to zero or near-zero levels is expected below the sediment-depths explored in this study, ushering acute carbon starvation for the chemoorganoheterotrophs *in situ*.

Under any fluctuating environmental condition adaptive responses of microbial communities determine their resilience and survival value (Lauro et al. 2009). In response to nutrient scarcity, an initial increase in cell number followed by a prolonged retention of the same is known to be a common survival strategy of bacteria (Amy and Morita 1983). In the biogeochemical context of the ASOMZ sediments, sensing the population level and maintaining a fixed cell density under starvation, in the same way as observed in the present isolates, could be a key quality ensuring survival of chemoorganoheterotrophs in the deeper horizons beyond 300 cmbsf. At the molecular level, signaling pathways involving autoinducer-2 are the main drivers of adaptation to carbon starvation and various other environmental stressors (McDougald et al. 2002), while the protein LuxS is involved in the secretion of autoinducer-2, the key sensor and messenger of information pertaining to cell density and nutrient concentration in the habitat. Therefore, the presence of several Lux regulon homologs in the genomes of the new isolates, as well as the metagenomes of the two cores investigated, could be further corroborative of a key role of quorum-sensing-based oligotrophy in sustaining the chemoorganoheterotrophic bacteria in the course of critical carbon depletion beyond the ∼3 m sediment horizon.

Notably, for all the current isolates, oligotrophic growth under aerobic or microaerobic (but not anaerobic) condition started with 10^3^ (but not 10^6^) CFU mL^-1^ of the culture medium, and stopped when a cell density of around 10^4^ or 10^5^ CFU mL^-1^ was reached after 96 hours of incubation; thereafter, the cell density remained unchanged even after 28 days of incubation (the cultures were not monitored for their CFU counts beyond 28 days; Table S9). Concurrently, though there was no growth for any of the nine isolates under the same conditions when incubation was started with a cell density of 10^6^ CFU mL^-1^, all the cultures retained around 10^4^ or 10^5^ CFU mL^-1^ from the 8^th^ day onward, till the 28^th^ day of incubation, when cell densities were last measured (Table S9). While similar responses to nutrient deprivation are common among microorganisms living in other oligotrophic ecosystems (Nimonkar et al. 2022), for the inhabitants of carbon-impoverished marine deep subsurfaces, *in situ* metabolism, and thereby community size, is said to be balanced against the ambient flux of carbon and energy, with cells recalibrating their metabolic rates down to such bare minimum levels that are essential for system maintenance, survival, and long-term viability (Jørgensen, 2011). In the context of the ASOMZ sediments, however, it is noteworthy that extreme oligotrophy in all the nine isolates was critically affected by the availability of O_2_ in the environment. For instance, during microaerobic growth, all the cultures had longer generation times, accompanied by delayed attainment of the steady cell density of 10^4^ or 10^5^ CFU mL^-1^, compared with the corresponding phenotypes exhibited under aerobic condition(Fig. 4, Table S9). Furthermore, anaerobic incubation in oligotrophic (as well as copiotrophic) medium, for all the ASOMZ-sediment isolates, irrespective of the starting cell density, led to the retention of zero CFU after 8-28 days. Thus persistence as non-growing viable cells could be a key mechanism for survival in the face of carbon starvation under aerobic to microaerobic conditions, but not under complete anoxia.

### Nitrate/nitrite reduction, and fermentation, as potential means of coping anoxia

Two out the nine ASOMZ strains characterized in this study, namely *Brucella* sp. JSBI001 and *H*. *bauzanensis* SMJS2, were found to be facultatively anaerobic with abilities to grow on yeast extract or acetate via nitrate and/or nitrite reduction in the absence of O_2_. While the genomic contents of these two strains corroborated their nitrate-/nitrite-respiring phenotypes, three more new isolates, namely *Halomonas* sp. BDJS001, *M*. *segnicrescens* AN1 and *Mesobacillus* sp. SBJS01, despite exhibiting no such phenotype in chemoorganoheterotrophic medium based on yeast extract or acetate, did possess all the genes necessary for respiring nitrate/nitrite to dinitrogen, ammonia or nitrous oxide. Therefore, possibilities remain that the nitrate/nitrite reduction capabilities of these organisms are coupled with some hitherto unknown catabolic processes that utilize substrates other than those which were tested in the present study. In such a scenario too, there remain four ASOMZ-sediment isolates (*Brevibacterium* strains BDJS002 and JSBI002, *G*. *hongkongensis* SMJS1, and *Stenotrophomonas* sp. SBJS02) whose survival in this sulfidic (anoxic) habitat is unexplainable by the biogeochemical data available for the ecosystem.

In the above context it is noteworthy that all the new strains isolated from the ASOMZ sediments could maintain a fraction of their cell population over prolonged anaerobic incubation by means of fermentative metabolism (Fig. 2, Table S7). Obligately aerobic bacteria, across diverse taxonomic groups and ecological systems, are known to withstand O_2_ limitation by using various fermentative metabolisms, often in conjunction with nitrate reduction (Eschbach et al. 2004; Van Keulen et al. 2007; Pinchuk et al. 2011; Berney et al. 2014; Falke et al. 2019; Gillett et al. 2023). It is, therefore, not unlikely thatnitrate reduction and fermentation collectively play a significant role in the sustenance of aerobic bacteria within the sulfidic (anoxic) sediments of the ASOMZ. However, further studies of pure-culture, as well as culture-independent, microbiology are required to establish the exact scopes of these metabolic processes *in situ*. Such information can subsequently be used to determine the actual biogeochemical implications that these processes potentially hold for *in situ* carbon cycling.

### NO dismutation, and other potential sources, of cryptic O_2_ for aerobic bacteria in OMZ sediments

Albeit the present state of knowledge does not offer any empirical insight to the absence or presence of O_2_ (at whatever miniscule concentration that may be) in the ASOMZ sediments, all information available thus far on the biogeochemistry of the system (Fernandes et al. 2018, 2022) indicate that this sulfidic habitat is essentially anoxic below a few centimeters from the seafloor (Bhattacharya et al. 2019). From a biochemical perspective it is noteworthy that aerobic bacteria can thrive on nanomolar, or even lower, O_2_ concentrations *in vitro* (Stolper et al. 2010), as well as in natural habitats (Ruff et al. 2023) including marine territories (Ulloa et al. 2012; Wright et al. 2012; Kalvelage et al. 2015; Garcia-Robledo et al. 2017) that are apparently anoxic. Whereas, “dark oxygen” produced by means of dismutation reactions is thought to provide potential sustenance to aerobic microbial communities in acutely O_2_-limited ancient groundwater ecosystems (Ruff et al. 2023), minute spatiotemporal variations in the O_2_ level of the chemical milieu have been hypothesized as the key driver of the stable co-occurrence of aerobic and anaerobic microorganisms in anoxic marine waters (Zakem et al. 2020). However, for the sedimentary realm of the ASOMZ, very little information is there about how the aerobic bacteria present could survive in this sulfidic habitat. As for the nearly anoxic water columns of global marine OMZs, feasibility of active aerobic metabolism *in situ* has been attributed to local oxygenic photosynthesis (Tiano et al. 2014; Garcia-Robledo et al. 2017), and prompt response of community metabolism to brisk spatiotemporal fluctuations in the availability of O_2_ (Dalsgaard et al. 2014). Furthermore, biogenic O_2_ produced via dismutation of chlorite has been hypothesized, based on metagenomic footprints and results of co-culture experiments, to sustain aerobic microorganisms in anoxic/hypoxic aquifers (Ruff et al. 2023), as well as the same ASOMZ sediment system (Bhattacharya et al. 2020) on which the present study was based. Notably, however, no oxyanion of chlorine (perchlorate, chlorate, or chlorite) was detected in either of the two ecosystems.

NO dismutation is a potential source of biogenic O_2_ in deep subsurface microbiomes (Ettwig et al. 2010; Kraft et al. 2022; Ruff et al. 2023). NO dismutases of archaea (Kraft et al. 2022) and bacteria (Ettwig et al. 2010) apparently convert NO to N_2_ and O_2_, with or without forming N_2_O as an intermediate respectively. Notably, in both the processes the substrate NO is derived from the reduction of nitrite (NO_2_), which in turn is an intermediate of denitrification (NO_3_ reduction) in bacteria, and a derivative of aerobic NH_3_ oxidation in archaea. In biogeochemical context of SSK42/5 and SSK42/6, the NO dismutation mechanism encountered in aerobic NH_3_-oxidizing archaea (such as *N*. *maritimus*) seems to be more feasible *in situ*, as the sediments are redox-stratified and largely unbioturbated, so NO_3_ is typically restricted to the surface and near-surface territories, whereas NH_3_ prevails at increasingly high concentrations along the sediment-depths due to extensive organic matter remineralization (Fernandes et al. 2018). Indeed, along SSK42/5 and SSK42/6, the presence of *Nitrosopumilus* species including *N*. *maritimus* (Fig. 7, Table S19), and that of nitric oxide dismutase (*nod*) gene homologs as well (Table S20), were corroborated by CDSs annotated within the assembled metagenomes. The occurrence of the genus in this sediment system was further evidenced by the analysis of PCR-amplified 16S rRNA gene sequences (Table S21). Incidentally, we did not detect any NO dismutase encoding sequence, or for that matter any CDS ascribable to *Nitrosopumilus* species, in the assembled and annotated metatranscriptome of SSK42/5 and SSK42/6. This could be reflective of the low coverage breadth of the current metatranscriptomic sequence data with respect to the functional microbiome of the ASOMZ sediments, and/or the low expressivity of the *in situ* populations of *Nitrosopumilus*.

*Nitrosopumilus* and other ammonia-oxidizing archaea are widely distributed in the global ocean, and their habitats include seawaters and sediments having O_2_ concentration below the detection limit of current technologies (Karner et al. 2001; Könneke et al. 2005; Molina et al. 2010; Walker et al. 2010; Beman et al. 2012; Stewart et al. 2012; Sollai et al. 2019). This, in conjunction with the discovery of an NO dismutation phenotype in *Pseudomonas aeruginosa* that releases up to 20_µM O_2_ to the extracellular milieu (Lichtenberg et al. 2021), and the identification of nitric oxide dismutase genes in taxonomically diverse bacteria (Murali et al. 2022; Ruff et al. 2023), suggests that the molecular attributes of O_2_ generation could be widespread across the taxonomic as well as ecological spectrum. So far as NO dismutation by *N*. *maritimus* is concerned, upon onset of anoxia in laboratory cultures, aerobic ammonia oxidation generates N_2_ and O_2_ via NO, NO, and then N_2_O; subsequently, the O_2_ produced can not only act as the electron acceptor for restarting aerobic oxidation of ammonia but also accumulate in the cells’ external chemical milieu (Kraft et al. 2022). Therefore, whereas it is tempting to hypothesize that within the apparently anoxic (sulfidic) sedimentary environment of the OMZ sediments, potential close association with cells of *Nitrosopumilus* could be a viable means of O_2_ for other obligate aerobes present *in situ*, only future experiments involving anaerobic co-cultures can ascertain whether *Nitrosopumilus* strains can indeed produce and transmit such surplus O_2_ that can sustain other adjoining aerobic microorganisms.

### Potential role of the aerobic chemoorganoheterotrophs in the carbon cycle of ASOMZ sediments

The sediment horizon explored in the present study of pure culture, as well as omics-based, microbiology is situated off the west coast of India, and is impinged by the acutely hypoxic waters of the Arabian Sea OMZ. In this kind of marine biogeochemical systems, water column productivity and organic carbon deposition to the seafloor are both very high; abundant complex organic matter is delivered to the seabed (Cavan et al. 2017; Jessen et al. 2017) since low concentrations of dissolved O_2_ in the overlying waters limit their biodegradation prior to deposition (Revsbech et al. 2009; Ulloa et al. 2012; Garcia-Robledo et al. 2017). Subsequently, within the sedimentary realm, notwithstanding the virtual absence of O_2_ beneath only a few cmbsf (Bhattacharya et al. 2019), efficient remineralization of the deposited organic matter takes place through the first few meters of the sediment horizon (Fernandes et al. 2018). Furthermore, it is remarkable that the catabolic degradation of the complex carbon compounds buried in these sediments takes place hand in hand with anaerobic microbial metabolisms that are dependent on the utilization of simple fatty acids as carbon source (van der Weijden et al. 1999; Schulte et al. 2000; Seiter et al. 2005; Bowles et al. 2014; Cowie et al. 2014; Fernandes et al. 2018; Ruvalcaba et al. 2020; Bhattacharya et al. 2021). Exceptionally high activities of anaerobic life processes in the surface and near-surface layers of these sediment systems are attributed to low sedimentation rates affording high O_2_-exposure time for the degradation of organic carbon in the surficial sediments despite acute hypoxia in the benthic waters (Bhattacharya et al. 2021). So far as the sites SSK42/5 and SSK42/6 underlying the approximate midpoint of the vertical span of the ASOMZ are concerned, copious organic matter deposition takes place at the sediment surface (organic matter constituted around 4% of the total sediment weight in the first few 10s cmbsf), while steady depletion of the total organic carbon content to ≤ 1% of the total sediment weight happens within a few 100s cmbsf (Fernandes et al. 2018). This gradual depletion trend is reflective of the fact that considerable proportions of the total organic carbon deposited to the seabed from the water column is degraded through the diagenetically matured, older and deeper, sediment layers amid an accentuating scarcity of oxidants down the sediment-depth. It is specifically in this biogeochemical context that the chemoorganoheterotrophic bacteria revealed in this study (alongside their physiological comparators that are also present *in situ*) emerge as the key players of carbon cycling in the ASOMZ sediments. Their versatile abilities to (i) catabolize, and grow on, complex organic compounds of both marine and terrestrial origin via aerobic respiration at low (as well as high) O_2_ concentration, (ii) grow anaerobically by reducing nitrate and/or nitrite, (iii) preserve a fraction of the cell population via fermentative metabolism under anaerobic condition, and (iv) grow by means of extreme oligotrophy at low cell density amid low (as well as high) O_2_ concentration, make them fit for prolonged carbon remineralization across the sediment horizon by utilizing whatever small amount of oxidants is available in time and space.

## SUPPLEMENTARY DATA

Supplementary information and data are available online in the form of a Word file named Supplimentary_Information.docx, and an Excel file named Supplimentary_Dataset.xlsx.

## DATA AVAILABILITY

GenBank accession numbers for the 16S rRNA genes, genomes, and plasmids (if any) of the new isolates are as follows: *Brevibacterium* sp. BDJS002 / MCC 5171 - OQ780597, CP116392; *Brevibacterium* sp. JSBI002 / MCC 5172 - OQ780604, CP110341; *Brucella* sp. JSBI001 / MCC 5134 - OQ780596, CP110339, CP110340; *Gordonia hongkongensis* SMJS1 / MCC 5348 - OQ780429, CP123381, CP123382, CP123383; *Halomonas* sp. BDJS001 / MCC 5128 - OQ780651, CP110535; *Halopseudomonas bauzanensis* SMJS2 / MCC 5363 - OQ780586, CP123445, CP123446; *Marinobacter segnicrescens* AN1 / MCC 5129 - OQ780700, CP110342; *Mesobacillus* sp. SBJS01 - OQ780701, CP109811; *Stenotrophomonas* sp. SBJS02 / MCC 5130 - OQ780703, CP109812.

All nucleotide sequence data have been deposited in NCBI Sequence Read Archive (SRA) or GenBank under the BioProject accession number PRJNA309469: The complete circular genome or plasmid sequences of the pure culture strains isolated have the GenBank accession numbers CP109811, CP109812, CP110339 through CP110342, CP110535, CP116392, CP123381 through CP123383, CP123445, and CP123446.

The whole metagenome shotgun sequence datasets have the Run Accession numbers SRR3646127, SRR3646128, SRR3646129, SRR3646130, SRR3646131, SRR3646132, SRR3646144, SRR3646145, SRR3646147, SRR3646148, SRR3646150, SRR3646151, SRR3646152, SRR3646153, SRR3646155, SRR3646156, SRR3646157, SRR3646158, SRR3646160, SRR3646161, SRR3646162, SRR3646163, SRR3646164, SRR3646165; SRR3570036, SRR3570038, SRR3577067, SRR3577068, SRR3577070, SRR3577071, SRR3577073, SRR3577076, SRR3577078, SRR3577079, SRR3577081, SRR3577082, SRR3577086, SRR3577087, SRR3577090, SRR3577311, SRR3577337, SRR3577338, SRR3577341, SRR3577343, SRR3577344, SRR3577345, SRR3577347, SRR3577349, SRR3577350, and SRR3577351.

The whole metatranscriptome shotgun sequence datasets have the Run Accession numbers SRR7967962, SRR9596399, SRR7983762, SRR9597791, and SRR7992674.

The sequence datasets obtained via PCR amplification of the V4-V5 regions of *Nitrosopumilus*-specific 16S rRNA genes have the Run Accession numbers SRR24121604, SRR24121705, SRR24121602, SRR24121623, SRR24123082, SRR24122693, SRR24121664, SRR24121706, SRR24121641, SRR24121637, SRR24121644, SRR24121647, SRR24121642, SRR24122688, SRR24122690, SRR24121638, SRR24121640, SRR24121648, SRR24121663, SRR24121649, SRR24121645, SRR24123085, SRR24122686, SRR24121636, and SRR24121643.

## ACKNOWLEDGEMENTS

We express gratitude to the Director of CSIR-National Institute of Oceanography for enabling the research cruise SSK42. Support extended by the CSIR-NIO Ship Cell and crew members of SSK42 is recognized gratefully. J.S. and S.D. received fellowships from Council of Scientific and Industrial Research, Government of India (GoI). M.M. and S.C. received fellowships from the Department of Biotechnology, GoI. S.B., N.M. and S.N. got their fellowships from Bose Institute.

## AUTHOR CONTRIBUTIONS

W.G. conceived the study, designed the experiments, interpreted the results, and wrote the paper. J.S. planned and performed the experiments, analyzed and curated the data, and also composed the paper. A.M. led the SSK42 mission and all geochemical studies therein; A.M. and A.P. made intellectual contributions to the present paper. M.M. performed microbiological experiments, while S.B., S.D., S.C., N.M. and S.N. performed omics studies. All authors read and ratified the manuscript.

