## Supplementary material for "Extremely oligotrophic and complex carbon degrading microaerobic bacteria from Arabian Sea oxygen minimum zone sediments": Figures S1 to S4 and Table S1, Tables S5 to S9

**Running Title:**	Carbon cyclers from marine sediments underlying a hypoxic water column

**Contents**

**Supplementary Figures**

**Figure S1.** Histograms showing the percentage-wise distributions of the chromosome- or plasmid-borne CDSs of the ASOMZ-sediment isolates across the different functional categories for clusters of orthologous genes (COGs).

**Figure S2.** Heat map comparing the distributions of all chromosome- or plasmid-borne CDSs of the ASOMZ-sediment isolates across the different COG categories.

**Figure S3.** Presence or absence of genes (CDSs) for some of the key enzymes of the various pathways of benzoate catabolism within the annotated genomes of the ASOMZ-sediment isolates.

**Figure S4.** Presence or absence of genes (CDSs) for some of the key enzymes of the various pathways of benzoate catabolism within the metatranscriptomes assembled from the different sediment samples of SSK42/5 and SSK42/6.

**Supplementary Tables**

**(Tables S2 through S4, and S10 through S21, are all more than one page long, so they have been given as individual sheets of an Excel Workbook named Supplementary_Dataset)**

**Table S1.** Sequence read archive (SRA) accession numbers of the whole genome sequence datasets obtained for the new isolates using short as well as long read technologies.

**Table S2.** Key attributes of the complete whole genome sequences of the ASOMZ-sediment isolates.

**Table S3.** Summary of the metagenome analyses (*de novo* assembly and annotation) carried out for the individual sediment samples of SSK42/5 and SSK42/6.

**Table S4.** Summary of the metatranscriptome analyses (*de novo* assembly and annotation) carried out for a number of sediment samples from SSK42/5 and SSK42/6.

**Table S5.** Sequence read archive (SRA) accession numbers of the Ion S5 sequence datasets obtained via PCR amplification of the V4-V5 regions of all *Nitrosopumilus*-specific 16S rRNA genes present within the metagenomes of SSK42/5.

**Table S6.** Sequence read archive (SRA) accession numbers of the Ion S5 sequence datasets obtained via PCR amplification of the V4-V5 regions of all *Nitrosopumilus*-specific 16S rRNA genes present within the metagenomes of SSK42/6.

**Table S7.** Increase or decrease in the CFU count of the nine OMZ-sediment isolates upon incubation in different ASW-based media, under aerobic, microaerobic, and anaerobic conditions.

**Table S8.** Increase or decrease in the CFU count of the nine OMZ-sediment isolates upon aerobic, microaerobic, or anaerobic incubation in ASW-based media supplemented with different complex carbon compounds.

**Table S9.** Increase or decrease in the CFU count of the nine OMZ-sediment isolates upon aerobic, microaerobic, or anaerobic incubation in ASW-based media supplemented with extremely low concentration of yeast extract (ASW_ELY).

**Table S10.** Genes identified within the genomes of the ASOMZ-sediment isolates for the different mechanisms of aerobic respiration.

**Table S11.** Genes identified within the genomes of the ASOMZ-sediment isolates for the different reactions of anaerobic respiration by nitrate/nitrite reduction, and fermentation.

**Table S12.** Genes possessed by the ASOMZ-sediment isolates for the breakdown of different glycosidic bonds within diverse polymeric carbohydrates and glycoconjugates.

**Table S13.** Genes identified within the genomes of the ASOMZ-sediment isolates for the catabolism of aromatic (benzoate) compounds.

**Table S14.** Genes identified within the genomes of the ASOMZ-sediment isolates for quorum sensing, and oligotrophy or starvation survival.

**Table S15.** The percentages of metagenomic read from individual sediment-samples of SSK42/5 and SSK42/6 that matched with sequences from the genomes of the different isolates.

**Table S16.** The percentages of metatranscriptomic read from individual sediment-samples of SSK42/5 and SSK42/6 that matched with sequences from the genomes of the different isolates.

**Table S17.** Numbers and percentages of genes identified as associated with aerobic and microaerobic respiration, nitrate/nitrite respiration, fermentation, catabolism of polymeric carbohydrates / glycoconjugates and benzoate, oligotrophy / starvation endurance, and quorum sensing, within the metatranscriptomes assembled from the different sediment samples of SSK42/5 and SSK42/6.

**Table S18.** Numbers and percentages of gene homolog identified within the different assembled metagenomes of SSK42/5 as being ascribable to *Nitrosopumilus maritimus* and *Nitrosopumilus* *sediminis*, on the basis of amino acid sequence homology.

**Table S19.** Numbers and percentages of gene homolog identified within the different assembled metagenomes of SSK42/6 as being ascribable to *Nitrosopumilus* *maritimus* and *Nitrosopumilus* *sediminis*, on the basis of amino acid sequence homology.

**Table S20.** Gene homologs (CDSs) identified within the different assembled metagenomes of SSK42/5 and SSK42/6 for the nitric oxide dismutase (*nod*) gene via BlastX analysis against a subject database that encompassed all known *nod* gene sequences.

**Table S21.** RDP classification of the Ion S5 reads that were obtained after sequencing of the PCR products generated from the different metagenomes of SSK42/5 and SSK42/6 using putatively Nitrosopumilus-specific 16S rRNA gene sequence primers.

**Supplementary Figures**

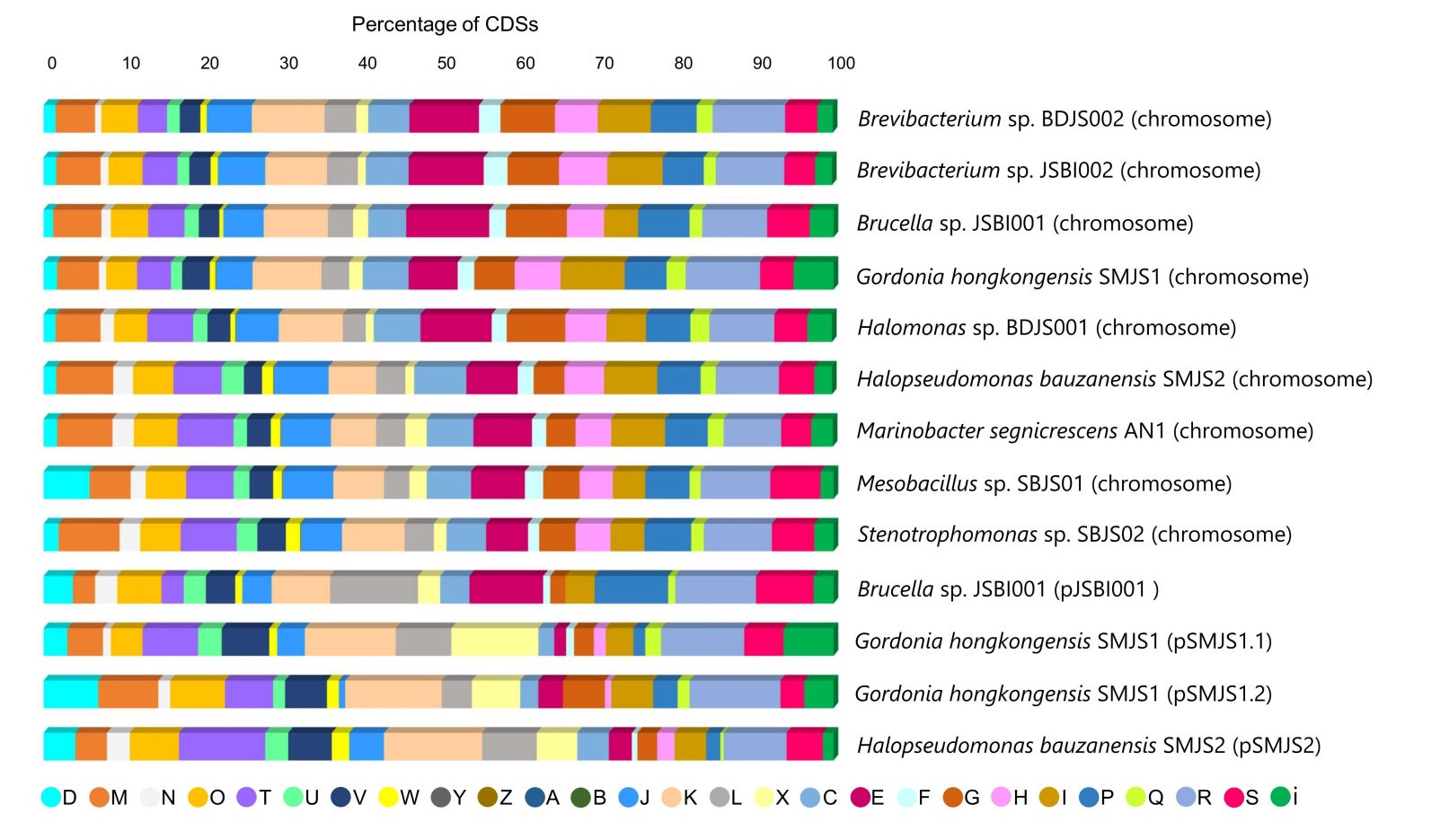

**Figure S1.** Histograms showing the percentage-wise distributions of the chromosome- or plasmid-borne CDSs of the ASOMZ-sediment isolates across the different functional categories for clusters of orthologous genes (COGs). The one-letter abbreviations indicated in the color code for the different functional categories stand for (A) RNA processing and modification; (B) Chromatin structure and dynamics; (C) Energy production and conversion; (D) Cell division and chromosome partitioning; (E) Amino acid metabolism and transport; (F) Nucleotide metabolism and transport; (G) Carbohydrate metabolism and transport; (H) Coenzyme metabolism; (I) Lipid metabolism; (J) Translation, including ribosome structure and biogenesis; (K) Transcription; (L) Replication, recombination and repair; (M) Cell wall structure and biogenesis and outer membrane; (N) Secretion, motility and chemotaxis; (O) Molecular chaperones and related functions; (P) Inorganic ion transport and metabolism; (Q) Secondary metabolites biosynthesis, transport and catabolism; (R) General functional prediction only; (S) No functional prediction; (T) Signal transduction; (V) Defense mechanisms; (W) Extracellular structures; (X) Mobilome: prophages, transposons; (Y) Nuclear structure; (Z) Cytoskeleton; (i) Genes which encode either hypothetical proteins or putatively functional proteins that do not belong to any COG.

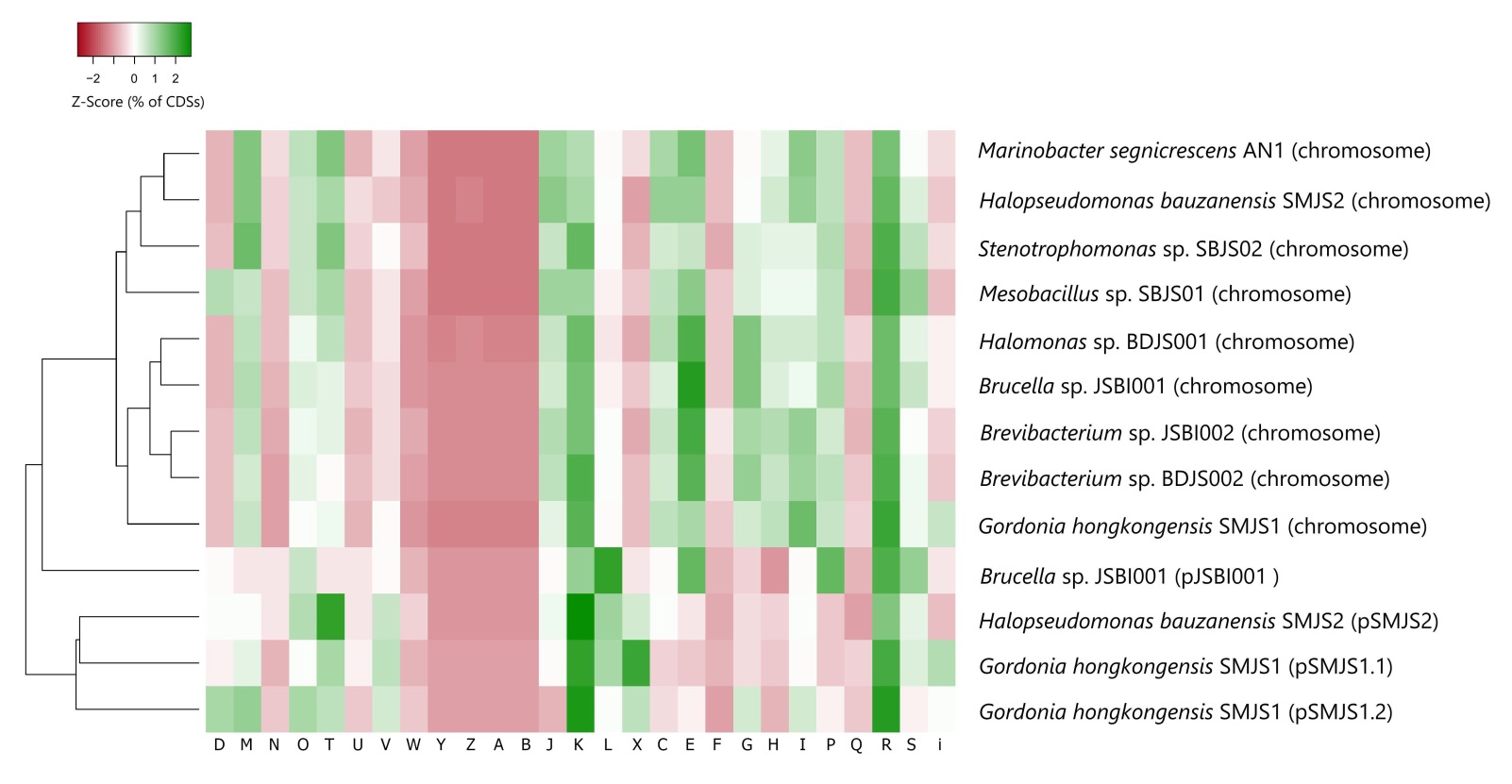

**Figure S2.** Heat map comparing the distributions of all chromosome- or plasmid-borne CDSs of the ASOMZ-sediment isolates across the different COG categories. For a given chromosome or plasmid, the percentage of its total CDSs ascribed to the different COG categories were first normalized by calculating the geometric means of the percentage values, and then dividing each percentage value by that mean (Table S2). Z-score normalization was performed upon the normalized CDS percentages across the individual chromosomes or plasmids. Z-scores were computed on a row by row basis by subtracting the mean of the normalized percentage values from each normalized percentage value and then dividing the individual remainders by the standard deviation of the normalized percentage values. As the rows were Z-Score scaled, cell colors represented how CDS percentages varied for a COG category across the chromosomes/plasmids, with white representing a row Z-Score of 0, while increasing positive and negative values are represented by the intensifying shades of blue and red respectively. The nine genomes and the four plasmids were further clustered in a dendogram (via Euclidean distance measurement) based on the Z-scores of their CDS distribution across the different COG categories. The one-letter abbreviations indicated for the different functional categories of COGs are same as those used in Figure S1.

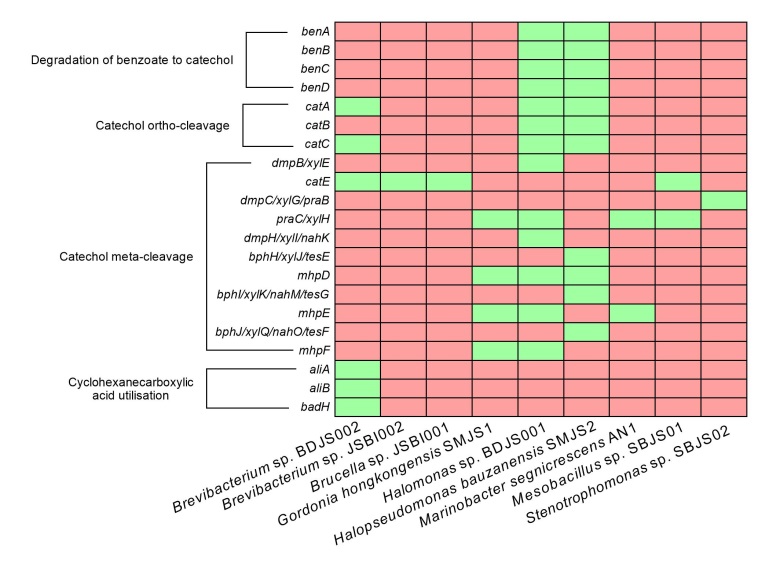

**Figure S3.** Presence (indicated by green boxes) or absence (indicated by red boxes) of genes (CDSs) for some of the key enzymes of the various pathways of benzoate catabolism within the annotated genomes of the ASOMZ-sediment isolates.

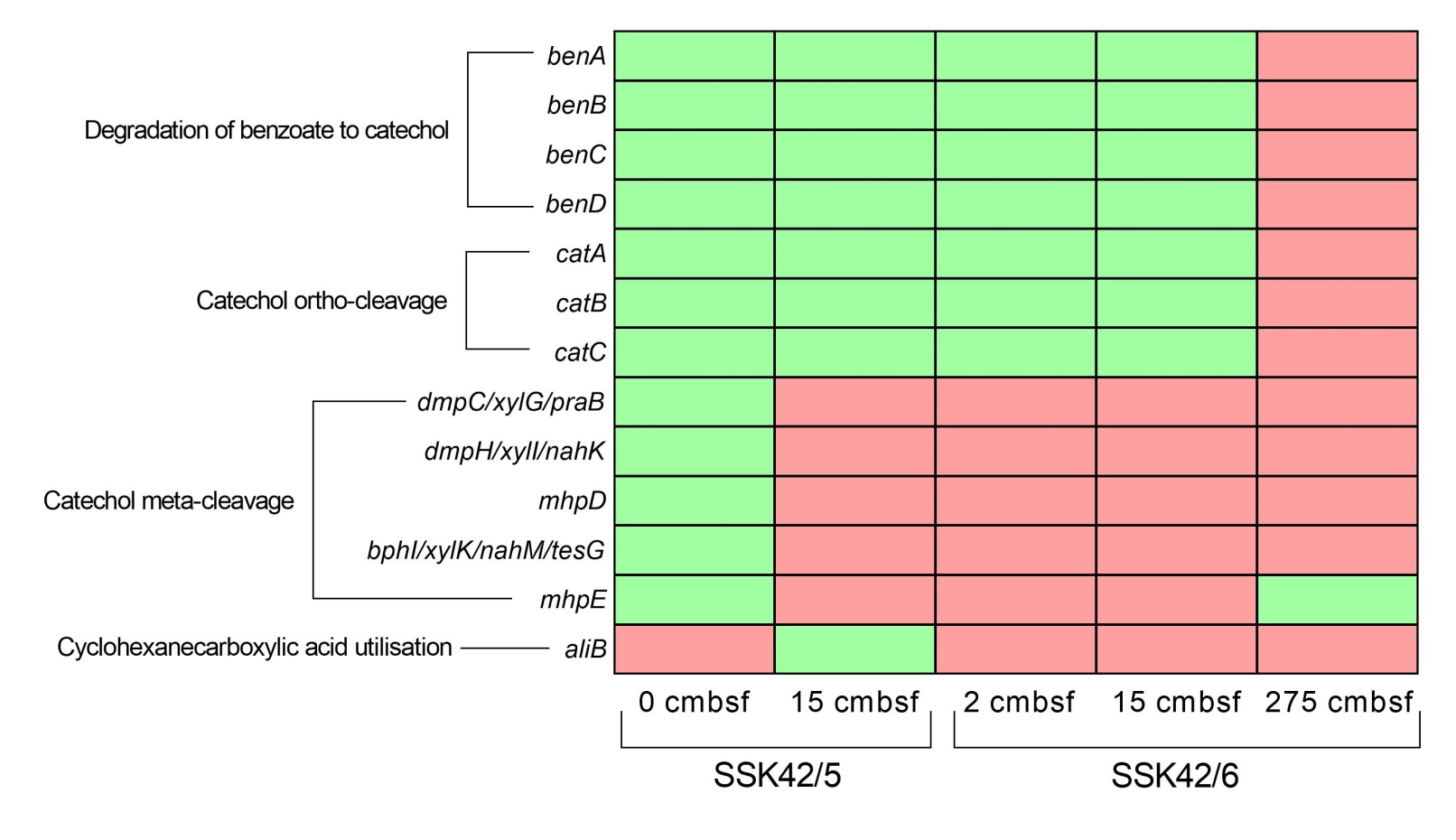

**Figure S4.** Presence (indicated by green boxes) or absence (indicated by red boxes) of genes (CDSs) for some of the key enzymes of the various pathways of benzoate catabolism within the metatranscriptomes assembled from the different sediment samples of SSK42/5 and SSK42/6.

**Supplementary Tables**

**Table S1.** Sequence read archive (SRA) accession numbers of the whole genome sequence datasets obtained for the new isolates using short as well as long read technologies.

| **Name of the isolate** | **BioSample accession number** | **Run accession number of the Ion S5 (short read) sequence dataset** | **Run accession number of the Oxford Nanopore (long read) sequence dataset** |
| --- | --- | --- | --- |
| *Brevibacterium* sp. BDJS002 | SAMN32648651 | SRR23097118 | SRR23097117 |
| *Brevibacterium* sp. JSBI002 | SAMN22418679 | SRR16493691 | SRR21903191 |
| *Brucella* sp. JSBI001 | SAMN22417814 | SRR16493532 | SRR21903039 |
| *Gordonia hongkongensis*  SMJS1 | SAMN34045557 | SRR24093915 | SRR24093914 |
| *Halomonas* sp. BDJS001 | SAMN31497625 | SRR22101745 | SRR22101744 |
| *Halopseudomonas*  *bauzanensis* SMJS2 | SAMN34065700 | SRR24060529 | SRR24060528 |
| *Marinobacter* *segnicrescens* AN1 | SAMN22405812 | SRR16481926 | SRR24138404 |
| *Mesobacillus* sp. SBJS01 | SAMN12273885 | SRR9698152 | SRR21902197 |
| *Stenotrophomonas* sp. SBJS02 | SAMN12369121 | SRR9843195 | SRR21902393 |

**Table S5.** Sequence read archive (SRA) accession numbers of the Ion S5 sequence datasets obtained via PCR amplification of the V4-V5 regions of all *Nitrosopumilus*-specific 16S rRNA genes present within the metagenomes of SSK42/5.

| **Sediment-depth explored (cmbsf)** | **BioSample accession number** | **Run accession number** |
| --- | --- | --- |
| 0 | SAMN34121403 | SRR24121604 |
| 15 | SAMN34121425 | SRR24121705 |
| 45 | SAMN34121439 | SRR24121602 |
| 60 | SAMN34121440 | SRR24121623 |
| 90 | SAMN34123134 | SRR24123082 |
| 120 | SAMN34123223 | SRR24122693 |
| 140 | SAMN34123238 | SRR24121664 |
| 160 | SAMN34123245 | SRR24121706 |
| 190 | SAMN34123376 | SRR24121641 |
| 220 | SAMN34123398 | SRR24121637 |
| 260 | SAMN34123501 | SRR24121644 |
| 295 | SAMN34123502 | SRR24121647 |

**Table S6.** Sequence read archive (SRA) accession numbers of the Ion S5 sequence datasets obtained via PCR amplification of the V4-V5 regions of all *Nitrosopumilus*-specific 16S rRNA genes present within the metagenomes of SSK42/6.

| **Sediment-depth explored (cmbsf)** | **BioSample accession number** | **Run accession number** |
| --- | --- | --- |
| 2 | SAMN34125098 | SRR24121642 |
| 30 | SAMN34125660 | SRR24122688 |
| 45 | SAMN34125692 | SRR24122690 |
| 60 | SAMN34125709 | SRR24121638 |
| 75 | SAMN34126542 | SRR24121640 |
| 90 | SAMN34126560 | SRR24121648 |
| 120 | SAMN34126586 | SRR24121663 |
| 135 | SAMN34126587 | SRR24121649 |
| 175 | SAMN34126588 | SRR24121645 |
| 220 | SAMN34126598 | SRR24123085 |
| 250 | SAMN34126641 | SRR24122686 |
| 265 | SAMN34126800 | SRR24121636 |
| 275 | SAMN34126802 | SRR24121643 |

**Table S7.** Increase or decrease in the CFU count of the nine OMZ-sediment isolates upon incubation in different ASW-based media, under aerobic, microaerobic, and anaerobic conditions.

| Culture medium | Days of incubation | *Brevibacterium* sp. BDJS002 | *Brevibacterium* sp. JSBI002 | *Brucella* sp. JSBI001 | *Gordonia hongkongensis*  SMJS1 | *Halomonas* sp. BDJS001 | *Halopseudomonas*  *bauzanensis* SMJS2 | *Marinobacter* *segnicrescens*  AN1 | *Mesobacillus* sp. SBJS01 | *Stenotrophomonas* sp. SBJS02 |
| --- | --- | --- | --- | --- | --- | --- | --- | --- | --- | --- |
| **Aerobic incubations** | | | | | | | | | | |
| ASW_HY | 0 | 3.1 × 10^6^ | 2.3 × 10^6^ | 3.7 × 10^6^ | 3.1 × 10^6^ | 1.3 × 10^6^ | 1.5 × 10^6^ | 4.8 × 10^6^ | 1.3 × 10^6^ | 1.4 × 10^6^ |
|  | 1 | 5.1 × 10^7^ | 2.7 × 10^8^ | 5.4 × 10^8^ | 2.3 × 10^7^ | 2.9 × 10^7^ | 3.1 × 10^7^ | 5.7 × 10^7^ | 3.2 × 10^8^ | 2.5 × 10^8^ |
|  | 2 | 3.2 × 10^8^ | - | - | 3.8 × 10^8^ | 5.1 × 10^8^ | 2.8 × 10^8^ | 3.1 × 10^8^ | - | - |
| ASW_A | 0 | 3.7 × 10^6^ | 3.7 × 10^6^ | 2.9 × 10^6^ | 1.5 × 10^6^ | 2.9 × 10^6^ | 4.1 × 10^6^ | 4.5 × 10^6^ | 3.7 × 10^6^ | 4.7 × 10^6^ |
|  | 1 | 4.9 × 10^7^ | 2.6 × 10^8^ | 1.9 × 10^8^ | 2.3 × 10^7^ | 3.7 × 10^7^ | 2.7 × 10^7^ | 2.2 × 10^7^ | 2.5 × 10^8^ | 2.3 × 10^8^ |
|  | 2 | 2.1 × 10^8^ | - | - | 2.6 × 10^8^ | 3.9 × 10^8^ | 3.8 × 10^8^ | 3.4 × 10^8^ | - | - |
| **Microaerobic incubation** | | | | | | | | | | |
| ASW_HY | 0 | 2.7 × 10^6^ | 3.9 × 10^6^ | 1.8 × 10^6^ | 2.5 × 10^6^ | 2.6 × 10^6^ | 3.7 × 10^6^ | 2.9 × 10^6^ | 3.9 × 10^6^ | 4.1 × 10^6^ |
|  | 4 | 3.1 × 10^7^ | 5.8 × 10^7^ | 6.7 × 10^7^ | 3.1 × 10^7^ | 1.9 × 10^7^ | 2.8 × 10^7^ | 3.9 × 10^7^ | 5.1 × 10^7^ | 6.2 × 10^7^ |
|  | 8 | 8.1 × 10^7^ | 2.7 × 10^8^ | 1.3 × 10^8^ | 9.3 × 10^7^ | 5.1 × 10^8^ | 1.9 × 10^8^ | 1.3 × 10^8^ | 3.5 × 10^8^ | 3.7 × 10^8^ |
| **Anaerobic incubations** | | | | | | | | | | |
| ASW_HY with seven electron acceptors* | 0 | 1.7 × 10^6^ | 3.2 × 10^6^ | 1.1 × 10^6^ | 5.2 × 10^6^ | 1.3 × 10^6^ | 2.3 × 10^6^ | 1.2 × 10^6^ | 1.3 × 10^6^ | 4.0 × 10^6^ |
|  | 4 | 2.8 × 10^3^ | 2.1 × 10^5^ | 2.6 × 10^6^ | 5.8 × 10^2^ | 3.2 × 10^3^ | 3.5 × 10^3^ | 1.1 × 10^5^ | 5.3 × 10^4^ | 2.9 × 10^3^ |
|  | 8 | 0 | 0 | 6.2 × 10^7^ | 0 | 0 | 0 | 0 | 0 | 0 |
|  | 12 | - | - | 9.1 × 10^8^ | - | - | - | - | - | - |
| ASW_HY  with NaNO_3_ | 0 | 1.9 × 10^6^ | 3.6 × 10^6^ | 1.9 × 10^6^ | 3.2 × 10^6^ | 1.7 × 10^6^ | 3.9 × 10^6^ | 1.7 × 10^6^ | 1.5 × 10^6^ | 4.3 × 10^6^ |
|  | 4 | 3.7 × 10^3^ | 2.8 × 10^4^ | 3.1 × 10^6^ | 6.1 × 10^2^ | 2.7 × 10^3^ | 3.3 × 10^3^ | 6.7 × 10^4^ | 1.9 × 10^4^ | 3.9 × 10^3^ |
|  | 8 | 0 | 0 | 4.3 × 10^7^ | 0 | 0 | 0 | 0 | 0 | 0 |
|  | 12 | - | - | 2.1 × 10^8^ | - | - | - | - | - | - |
| ASW_HY  with NaNO_2_ | 0 | 1.8 × 10^6^ | 4.2 × 10^6^ | 2.8 × 10^6^ | 4.3 × 10^6^ | 4.2 × 10^6^ | 2.2 × 10^6^ | 2.6 × 10^6^ | 1.5 × 10^6^ | 2.7 × 10^6^ |
|  | 4 | 2.8 × 10^2^ | 1.5 × 10^3^ | 5.3 × 10^6^ | 2.1 × 10^2^ | 2.8 × 10^2^ | 1.8 × 10^2^ | 3.8 × 10^2^ | 1.2 × 10^3^ | 4.1 × 10^2^ |
|  | 8 | 0 | 0 | 2.3 × 10^7^ | 0 | 0 | 0 | 0 | 0 | 0 |
|  | 12 | - | - | 2.6 × 10^8^ | - | - | - | - | - | - |
| ASW_HY  with Fe_2_O_3_ | 0 | 3.7 × 10^6^ | 3.2 × 10^6^ | 2.1 × 10^6^ | 5.2 × 10^6^ | 1.3 × 10^6^ | 2.3 × 10^6^ | 1.9 × 10^6^ | 1.3 × 10^6^ | 4.3 × 10^6^ |
|  | 4 | 2.1 × 10^3^ | 1.9 × 10^4^ | 1.9 × 10^4^ | 2.8 × 10^2^ | 3.1 × 10^3^ | 3.1 × 10^2^ | 3.2 × 10^4^ | 3.5 × 10^3^ | 2.4 × 10^3^ |
|  | 8 | 0 | 0 | 0 | 0 | 0 | 0 | 0 | 0 | 0 |
| ASW_HY  with MnO_2_ | 0 | 3.4 × 10^6^ | 4.9 × 10^6^ | 3.1 × 10^6^ | 3.2 × 10^6^ | 1.7 × 10^6^ | 3.5 × 10^6^ | 1.7 × 10^6^ | 2.5 × 10^6^ | 3.5 × 10^6^ |
|  | 4 | 2.1 × 10^3^ | 2.1 × 10^5^ | 2.9 × 10^4^ | 1.8 × 10^2^ | 3.7 × 10^3^ | 4.1 × 10^2^ | 3.2 × 10^3^ | 1.9 × 10^4^ | 2.9 × 10^3^ |
|  | 8 | 0 | 0 | 0 | 0 | 0 | 0 | 0 | 0 | 0 |
| ASW_HY  with Na_2_SO_4_ | 0 | 2.5 × 10^6^ | 4.5 × 10^6^ | 2.9 × 10^6^ | 4.2 × 10^6^ | 3.7 × 10^6^ | 3.9 × 10^6^ | 2.5 × 10^6^ | 2.7 × 10^6^ | 4.5 × 10^6^ |
|  | 4 | 2.1 × 10^3^ | 2.1 × 10^5^ | 1.7 × 10^4^ | 2.6 × 10^2^ | 1.5 × 10^3^ | 3.1 × 10^2^ | 3.3 × 10^3^ | 3.9 × 10^4^ | 1.9 × 10^3^ |
|  | 8 | 0 | 0 | 0 | 0 | 0 | 0 | 0 | 0 | 0 |
| ASW_HY  with (CH_3_)_2_SO | 0 | 3.5 × 10^6^ | 2.9 × 10^6^ | 2.1 × 10^6^ | 2.2 × 10^6^ | 3.5 × 10^6^ | 3.3 × 10^6^ | 2.3 × 10^6^ | 3.5 × 10^6^ | 2.9 × 10^6^ |
|  | 4 | 2.1 × 10^3^ | 2.1 × 10^5^ | 2.9 × 10^4^ | 3.8 × 10^2^ | 2.7 × 10^3^ | 2.1 × 10^2^ | 3.9 × 10^3^ | 1.6 × 10^4^ | 1.3 × 10^3^ |
|  | 8 | 0 | 0 | 0 | 0 | 0 | 0 | 0 | 0 | 0 |
| ASW_HY  with (CH_3_)_3_NO | 0 | 3.2 × 10^6^ | 4.2 × 10^6^ | 3.3 × 10^6^ | 4.2 × 10^6^ | 3.7 × 10^6^ | 4.3 × 10^6^ | 2.7 × 10^6^ | 1.5 × 10^6^ | 3.5 × 10^6^ |
|  | 4 | 2.1 × 10^3^ | 3.1 × 10^5^ | 1.9 × 10^4^ | 1.7 × 10^2^ | 1.7 × 10^3^ | 3.1 × 10^2^ | 3.1 × 10^3^ | 1.7 × 10^4^ | 2.7 × 10^3^ |
|  | 8 | 0 | 0 | 0 | 0 | 0 | 0 | 0 | 0 | 0 |
| ASW_HY  with humic acids and Fe_2_O_3_ | 0 | 3.7 × 10^6^ | 6.2 × 10^6^ | 3.5 × 10^6^ | 1.9 × 10^6^ | 1.3 × 10^6^ | 2.4 × 10^6^ | 4.2 × 10^6^ | 1.3 × 10^6^ | 4.5 × 10^6^ |
|  | 4 | 5.8 × 10^3^ | 2.9 × 10^5^ | 2.8 × 10^5^ | 1.8 × 10^2^ | 4.7 × 10^3^ | 3.9 × 10^3^ | 5.1 × 10^5^ | 3.3 × 10^4^ | 7.9 × 10^3^ |
|  | 8 | 0 | 0 | 0 | 0 | 0 | 0 | 0 | 0 | 0 |
| ASW_A with seven electron acceptors* | 0 | 2.7 × 10^6^ | 4.2 × 10^6^ | 5.3 × 10^6^ | 4.2 × 10^6^ | 3.3 × 10^6^ | 3.7 × 10^6^ | 3.2 × 10^6^ | 3.1 × 10^6^ | 3.5 × 10^6^ |
|  | 4 | 6.8 × 10^3^ | 3.9 × 10^4^ | 3.8 × 10^3^ | 3.6 × 10^2^ | 4.7 × 10^3^ | 8.5 × 10^6^ | 2.1 × 10^2^ | 2.3 × 10^3^ | 2.3 × 10^2^ |
|  | 8 | 0 | 0 | 0 | 0 | 0 | 2.1 × 10^7^ | 0 | 0 | 0 |
|  | 12 | - | - | - | - | - | 7.7 × 10^8^ | - | - | - |
| ASW_A with NaNO_3_ | 0 | 1.1 × 10^6^ | 1.6 × 10^6^ | 3.9 × 10^6^ | 3.1 × 10^6^ | 4.1 × 10^6^ | 2.4 × 10^6^ | 2.7 × 10^6^ | 1.9 × 10^6^ | 2.8 × 10^6^ |
|  | 4 | 2.1 × 10^3^ | 2.9 × 10^2^ | 3.1 × 10^3^ | 3.8 × 10^2^ | 3.9 × 10^2^ | 2.9 × 10^2^ | 3.9 × 10^1^ | 2.8 × 10^3^ | 1.8 × 10^2^ |
|  | 8 | 0 | 0 | 0 | 0 | 0 | 0 | 0 | 0 | 0 |
| ASW_A with NaNO_2_ | 0 | 1.5 × 10^6^ | 4.5 × 10^6^ | 2.9 × 10^6^ | 4.5 × 10^6^ | 4.3 × 10^6^ | 2.8 × 10^6^ | 2.2 × 10^6^ | 1.4 × 10^6^ | 2.6 × 10^6^ |
|  | 4 | 3.1 × 10^3^ | 2.5 × 10^2^ | 6.1 × 10^3^ | 1.8 × 10^2^ | 3.5 × 10^3^ | 7.5 × 10^6^ | 4.1 × 10^2^ | 1.9 × 10^3^ | 2.3 × 10^2^ |
|  | 8 | 0 | 0 | 0 | 0 | 0 | 1.9 × 10^7^ | 0 | 0 | 0 |
|  | 12 | - | - | - | - | - | 5.6 × 10^8^ | - | - | - |
| ASW_A with Fe_2_O_3_ | 0 | 1.9 × 10^6^ | 4.2 × 10^6^ | 5.1 × 10^6^ | 3.2 × 10^6^ | 3.5 × 10^6^ | 2.4 × 10^6^ | 3.2 × 10^6^ | 2.1 × 10^6^ | 3.8 × 10^6^ |
|  | 4 | 5.1 × 10^2^ | 2.9 × 10^4^ | 2.8 × 10^3^ | 2.6 × 10^2^ | 1.7 × 10^2^ | 1.9 × 10^2^ | 2.7 × 10^2^ | 4.3 × 10^3^ | 2.2 × 10^2^ |
|  | 8 | 0 | 0 | 0 | 0 | 0 | 0 | 0 | 0 | 0 |
| ASW_A with MnO_2_ | 0 | 2.9 × 10^6^ | 3.2 × 10^6^ | 3.1 × 10^6^ | 4.2 × 10^6^ | 3.3 × 10^6^ | 3.4 × 10^6^ | 2.2 × 10^6^ | 3.7 × 10^6^ | 3.5 × 10^6^ |
|  | 4 | 5.8 × 10^2^ | 2.7 × 10^4^ | 1.8 × 10^3^ | 1.6 × 10^2^ | 4.7 × 10^2^ | 1.9 × 10^2^ | 3.9 × 10^2^ | 2.5 × 10^3^ | 2.8 × 10^2^ |
|  | 8 | 0 | 0 | 0 | 0 | 0 | 0 | 0 | 0 | 0 |
| ASW_A with Na_2_SO_4_ | 0 | 1.9 × 10^6^ | 4.1 × 10^6^ | 4.1 × 10^6^ | 2.9 × 10^6^ | 3.5 × 10^6^ | 3.8 × 10^6^ | 3.7 × 10^6^ | 3.6 × 10^6^ | 3.9 × 10^6^ |
|  | 4 | 5.1 × 10^2^ | 3.5 × 10^4^ | 2.8 × 10^3^ | 3.6 × 10^2^ | 2.7 × 10^2^ | 1.7 × 10^2^ | 4.1 × 10^2^ | 2.8 × 10^3^ | 2.5 × 10^2^ |
|  | 8 | 0 | 0 | 0 | 0 | 0 | 0 | 0 | 0 | 0 |
| ASW_A with (CH_3_)_2_SO | 0 | 3.9 × 10^6^ | 4.3 × 10^6^ | 3.9 × 10^6^ | 3.2 × 10^6^ | 3.3 × 10^6^ | 3.7 × 10^6^ | 3.5 × 10^6^ | 3.7 × 10^6^ | 3.3 × 10^6^ |
|  | 4 | 5.1 × 10^2^ | 1.9 × 10^4^ | 1.8 × 10^3^ | 1.6 × 10^2^ | 3.7 × 10^2^ | 1.9 × 10^2^ | 2.9 × 10^2^ | 2.3 × 10^3^ | 2.1 × 10^2^ |
|  | 8 | 0 | 0 | 0 | 0 | 0 | 0 | 0 | 0 | 0 |
| ASW_A with (CH_3_)_3_NO | 0 | 2.7 × 10^6^ | 4.2 × 10^6^ | 5.1 × 10^6^ | 2.7 × 10^6^ | 3.3 × 10^6^ | 3.5 × 10^6^ | 3.5 × 10^6^ | 3.9 × 10^6^ | 3.1 × 10^6^ |
|  | 4 | 5.7 × 10^2^ | 2.9 × 10^4^ | 2.8 × 10^3^ | 1.6 × 10^2^ | 2.3 × 10^2^ | 1.4 × 10^2^ | 4.1 × 10^2^ | 2.6 × 10^3^ | 2.3 × 10^2^ |
|  | 8 | 0 | 0 | 0 | 0 | 0 | 0 | 0 | 0 | 0 |
| ASW_A with humic acids and Fe_2_O_3_ | 0 | 2.1 × 10^6^ | 2.2 × 10^6^ | 2.1 × 10^6^ | 3.2 × 10^6^ | 3.3 × 10^6^ | 3.4 × 10^6^ | 4.2 × 10^6^ | 3.1 × 10^6^ | 3.5 × 10^6^ |
|  | 4 | 4.7 × 10^2^ | 2.9 × 10^4^ | 1.8 × 10^3^ | 3.6 × 10^2^ | 1.7 × 10^2^ | 1.2 × 10^2^ | 2.3 × 10^2^ | 2.7 × 10^3^ | 2.7 × 10^2^ |
|  | 8 | 0 | 0 | 0 | 0 | 0 | 0 | 0 | 0 | 0 |
| ASW_Glucose | 0 | 3.3 × 10^6^ | 2.6 × 10^6^ | 7.7 × 10^6^ | 7.6 × 10^6^ | 5.5 × 10^6^ | 5.3 × 10^6^ | 1.8 × 10^6^ | 1.8 × 10^6^ | 9.0 × 10^6^ |
|  | 4 | 1.5 × 10^6^ | 2.5 × 10^6^ | 5.2 × 10^6^ | 4.1 × 10^6^ | 1.7 × 10^6^ | 4.1 × 10^6^ | 1.0 × 10^6^ | 1.1 × 10^6^ | 5.2 × 10^6^ |
|  | 8 | 5.0 × 10^5^ | 5.0 × 10^5^ | 3.7 × 10^6^ | 3.7 × 10^6^ | 2.2 × 10^6^ | 2.9 × 10^6^ | 3.0 × 10^5^ | 9.3 × 10^5^ | 3.1 × 10^6^ |
|  | 16 | 3.0 × 10^4^ | 4.0 × 10^5^ | 2.1 × 10^4^ | 7.0 × 10^5^ | 9.0 × 10^5^ | 2.5 × 10^6^ | 4.1 × 10^5^ | 9.3 × 10^3^ | 2.2 × 10^6^ |
|  | 32 | 6.0 × 10^3^ | 1.1 × 10^4^ | 1.5 × 10^4^ | 8.3 × 10^3^ | 5.9 × 10^4^ | 1.5 × 10^3^ | 7.1 × 10^4^ | 8.5 × 10^2^ | 1.5 × 10^5^ |
|  | 64 | 2.5 × 10^2^ | 2.1 × 10^3^ | 1.1 × 10^3^ | 2.1 × 10^1^ | 2.3 × 10^2^ | 2.5 × 10^2^ | 2.1 × 10^2^ | 6.7 × 10^2^ | 1.9 × 10^2^ |
| ASW_Pyruvate | 0 | 1.5 × 10^3^ | 4.6 × 10^6^ | 7.7 × 10^6^ | 7.6 × 10^6^ | 5.5 × 10^6^ | 5.3 × 10^6^ | 1.8 × 10^6^ | 1.8 × 10^6^ | 9.0 × 10^6^ |
|  | 4 | 2.6 × 10^6^ | 4.1 × 10^6^ | 5.7 × 10^6^ | 6.8 × 10^6^ | 3.9 × 10^6^ | 4.6 × 10^6^ | 1.2 × 10^6^ | 1.4 × 10^6^ | 7.9 × 10^6^ |
|  | 8 | 3.1 × 10^5^ | 3.7 × 10^6^ | 4.2 × 10^6^ | 6.5 × 10^6^ | 3.7 × 10^6^ | 3.6 × 10^6^ | 8.0 × 10^5^ | 1.3 × 10^6^ | 7.7 × 10^6^ |
|  | 16 | 2.5 × 10^4^ | 2.8 × 10^6^ | 3.2 × 10^4^ | 7.9 × 10^5^ | 9.0 × 10^4^ | 3.5 × 10^4^ | 4.9 × 10^4^ | 9.2 × 10^4^ | 2.3 × 10^4^ |
|  | 32 | 9.0 × 10^3^ | 4.3 × 10^4^ | 2.5 × 10^3^ | 3.1 × 10^3^ | 6.8 × 10^2^ | 1.8 × 10^3^ | 1.5 × 10^3^ | 8.4 × 10^2^ | 1.1 × 10^3^ |
|  | 64 | 2.3 × 10^1^ | 1.2 × 10^2^ | 2.0 × 10^2^ | 2.5 × 10^1^ | 3.5 × 10^1^ | 1.2 × 10^1^ | 3.7 × 10^1^ | 6.4 × 10^1^ | 3.1 × 10^1^ |
| ASW | 0 | 1.7 × 10^6^ | 3.2 × 10^6^ | 2.1 × 10^6^ | 5.2 × 10^6^ | 1.3 × 10^6^ | 2.3 × 10^6^ | 1.2 × 10^6^ | 1.3 × 10^6^ | 4.0 × 10^6^ |
|  | 4 | 2.1 × 10^4^ | 1.5 × 10^3^ | 1.2 × 10^5^ | 5.8 × 10^2^ | 3.2 × 10^3^ | 3.5 × 10^3^ | 2.9 × 10^1^ | 5.3 × 10^4^ | 2.9 × 10^3^ |
|  | 8 | 0 | 0 | 1.2 × 10^3^ | 0 | 0 | 0 | 0 | 0 | 0 |
|  | 16 | - | - | 0 | - | - | - | - | - | - |

* The seven terminal electron acceptors provided here as respiratory substrates were NaNO_3_, NaNO_2,_ Fe_2_O_3_, MnO_2_, Na_2_SO_4_, (CH_3_)_2_SO and (CH_3_)_3_NO.

**Table S8.** Increase or decrease in the CFU count of the nine OMZ-sediment isolates upon aerobic, microaerobic, or anaerobic incubation in ASW-based media supplemented with different complex carbon compounds.

| Incubation condition | Days of incubation | *Brevibacterium* sp. BDJS002 | *Brevibacterium* sp. JSBI002 | *Brucella* sp. JSBI001 | *Gordonia hongkongensis*  SMJS1 | *Halomonas* sp. BDJS001 | *Halopseudomonas*  *bauzanensis* SMJS2 | *Marinobacter* *segnicrescens*  AN1 | *Mesobacillus* sp. SBJS01 | *Stenotrophomonas* sp. SBJS02 |
| --- | --- | --- | --- | --- | --- | --- | --- | --- | --- | --- |
| **Culture medium used:** **ASW_Agar** | | | | | | | | | | |
| Aerobic | 0 | 1.8 × 10^6^ | 1.4 × 10^6^ | 2.5 × 10^6^ | 1.2 × 10^6^ | 5.6 × 10^6^ | 1.9 × 10^6^ | 1.3 × 10^6^ | 3.7 × 10^6^ | 4.8 × 10^6^ |
|  | 4 | 1.4 × 10^6^ | 1.1 × 10^8^ | 4.0 × 10^8^ | 5.4 × 10^5^ | 2.5 × 10^7^ | 3.5 × 10^5^ | 1.0 × 10^6^ | 2.5 × 10^8^ | 8.9 × 10^7^ |
| Microaerobic | 0 | 2.6 × 10^6^ | 1.6 × 10^6^ | 3.1 × 10^6^ | 5.7 × 10^6^ | 2.1 × 10^6^ | 1.2 × 10^6^ | 2.6 × 10^6^ | 1.2 × 10^6^ | 4.1 × 10^6^ |
|  | 4 | 7.4 × 10^5^ | 1.1 × 10^7^ | 9.3 × 10^6^ | 1.7 × 10^5^ | 1.6 × 10^7^ | 1.9 × 10^5^ | 2.6 × 10^5^ | 4.7 × 10^6^ | 1.3 × 10^6^ |
|  | 8 | 1.9 × 10^5^ | 3.9 × 10^7^ | 6.2 × 10^7^ | 8.2 × 10^4^ | 3.6 × 10^7^ | 4.7 × 10^4^ | 4.5 × 10^4^ | 1.1 × 10^7^ | 1.9 × 10^6^ |
| Anaerobic* | 0 | 1.6 × 10^6^ | 4.6 × 10^6^ | 2.5 × 10^6^ | 1.2 × 10^6^ | 5.6 × 10^6^ | 1.9 × 10^6^ | 1.3 × 10^6^ | 4.2 × 10^6^ | 4.8 × 10^6^ |
|  | 4 | 2.2 × 10^5^ | 2.3 × 10^4^ | 2.8 × 10^5^ | 5.4 × 10^4^ | 2.9 × 10^3^ | 4.2 × 10^5^ | 4.2 × 10^4^ | 7.4 × 10^4^ | 5.3 × 10^3^ |
| **Culture medium used: ASW_Alginate** | | | | | | | | | | |
| Aerobic | 0 | 1.8 × 10^6^ | 1.4 × 10^6^ | 2.5 × 10^6^ | 1.2 × 10^6^ | 1.3 × 10^6^ | 1.9 × 10^6^ | 1.8 × 10^6^ | 3.7 × 10^6^ | 4.8 × 10^6^ |
|  | 4 | 8.7 × 10^5^ | 1.3 × 10^8^ | 8.9 × 10^8^ | 4.8 × 10^7^ | 4.2 × 10^8^ | 1.8 × 10^5^ | 6.4 × 10^5^ | 5.9 × 10^7^ | 4.5 × 10^6^ |
| Microaerobic | 0 | 2.6 × 10^6^ | 1.6 × 10^6^ | 3.1 × 10^6^ | 5.7 × 10^6^ | 2.1 × 10^6^ | 1.2 × 10^6^ | 2.6 × 10^6^ | 1.2 × 10^6^ | 4.1 × 10^6^ |
|  | 4 | 5.4 × 10^5^ | 3.1 × 10^7^ | 4.6 × 10^6^ | 8.2 × 10^6^ | 1.5 × 10^7^ | 8.7 × 10^4^ | 1.8 × 10^5^ | 4.8 × 10^6^ | 1.2 × 10^6^ |
|  | 8 | 1.7 × 10^5^ | 3.3 × 10^7^ | 6.3 × 10^7^ | 3.0 × 10^7^ | 1.1 × 10^8^ | 1.7 × 10^4^ | 2.9 × 10^4^ | 9.7 × 10^6^ | 7.1 × 10^5^ |
| Anaerobic* | 0 | 1.6 × 10^6^ | 4.6 × 10^6^ | 4.4 × 10^6^ | 1.2 × 10^6^ | 5.6 × 10^6^ | 1.9 × 10^6^ | 1.3 × 10^6^ | 4.2 × 10^6^ | 5.1 × 10^6^ |
|  | 4 | 1.4 × 10^4^ | 1.6 × 10^4^ | 5.3 × 10^5^ | 1.2 × 10^4^ | 3.5 × 10^3^ | 2.3 × 10^5^ | 2.1 × 10^4^ | 1.1 × 10^4^ | 3.2 × 10^3^ |
| **Culture medium used: ASW_Carrageenan** | | | | | | | | | | |
| Aerobic | 0 | 1.8 × 10^6^ | 4.6 × 10^6^ | 4.4 × 10^6^ | 1.2 × 10^6^ | 5.6 × 10^6^ | 1.9 × 10^6^ | 1.8 × 10^6^ | 5.4 × 10^6^ | 2.5 × 10^6^ |
|  | 4 | 1.7 × 10^6^ | 4.1 × 10^7^ | 5.0 × 10^7^ | 2.4 × 10^5^ | 1.5 × 10^7^ | 2.5 × 10^4^ | 1.7 × 10^5^ | 9.2 × 10^6^ | 2.8 × 10^6^ |
| Microaerobic | 0 | 2.6 × 10^6^ | 1.6 × 10^6^ | 3.1 × 10^6^ | 5.7 × 10^6^ | 2.1 × 10^6^ | 1.2 × 10^6^ | 2.6 × 10^6^ | 1.2 × 10^6^ | 4.1 × 10^6^ |
|  | 4 | 4.3 × 10^5^ | 5.1 × 10^6^ | 6.1 × 10^6^ | 1.1 × 10^5^ | 4.4 × 10^6^ | 1.1 × 10^4^ | 5.9 × 10^4^ | 1.3 × 10^6^ | 2.7 × 10^6^ |
|  | 8 | 5.7 × 10^4^ | 2.4 × 10^7^ | 1.4 × 10^7^ | 4.3 × 10^4^ | 2.6 × 10^7^ | 5.8 × 10^3^ | 1.1 × 10^4^ | 3.0 × 10^6^ | 9.3 × 10^5^ |
| Anaerobic* | 0 | 1.6 × 10^6^ | 4.6 × 10^6^ | 4.4 × 10^6^ | 1.2 × 10^6^ | 5.6 × 10^6^ | 1.9 × 10^6^ | 1.5 × 10^6^ | 5.4 × 10^6^ | 1.4 × 10^6^ |
|  | 4 | 1.6 × 10^5^ | 5.8 × 10^4^ | 4.5 × 10^4^ | 4.5 × 10^5^ | 2.8 × 10^3^ | 7.4 × 10^4^ | 1.3 × 10^4^ | 4.1 × 10^4^ | 5.9 × 10^3^ |
| **Culture medium used: ASW_Chitosan** | | | | | | | | | | |
| Aerobic | 0 | 1.8 × 10^6^ | 4.6 × 10^6^ | 4.4 × 10^6^ | 1.2 × 10^6^ | 5.6 × 10^6^ | 1.9 × 10^6^ | 1.4 × 10^6^ | 5.4 × 10^6^ | 2.1 × 10^6^ |
|  | 4 | 7.9 × 10^5^ | 1.5 × 10^7^ | 3.3 × 10^7^ | 5.8 × 10^4^ | 5.4 × 10^6^ | 5.7 × 10^5^ | 8.0 × 10^5^ | 7.8 × 10^7^ | 1.1 × 10^7^ |
| Microaerobic | 0 | 2.6 × 10^6^ | 1.6 × 10^6^ | 3.1 × 10^6^ | 5.7 × 10^6^ | 2.1 × 10^6^ | 1.2 × 10^6^ | 2.6 × 10^6^ | 1.2 × 10^6^ | 4.1 × 10^6^ |
|  | 4 | 3.1 × 10^5^ | 5.3 × 10^5^ | 3.3 × 10^6^ | 1.1 × 10^4^ | 7.8 × 10^5^ | 2.3 × 10^5^ | 4.9 × 10^5^ | 5.8 × 10^6^ | 1.9 × 10^6^ |
|  | 8 | 1.4 × 10^5^ | 7.0 × 10^4^ | 1.1 × 10^7^ | 8.3 × 10^3^ | 9.5 × 10^4^ | 1.5 × 10^5^ | 1.2 × 10^5^ | 8.9 × 10^6^ | 3.2 × 10^6^ |
| Anaerobic* | 0 | 1.6 × 10^6^ | 4.6 × 10^6^ | 4.2 × 10^6^ | 1.2 × 10^6^ | 5.6 × 10^6^ | 1.9 × 10^6^ | 1.7 × 10^6^ | 5.4 × 10^6^ | 2.1 × 10^6^ |
|  | 4 | 3.9 × 10^4^ | 7.2 × 10^4^ | 2.3 × 10^5^ | 5.7 × 10^5^ | 7.1 × 10^3^ | 4.7 × 10^5^ | 2.4 × 10^4^ | 2.2 × 10^4^ | 4.3 × 10^3^ |
| **Culture medium used: ASW_Cellulose** | | | | | | | | | | |
| Aerobic | 0 | 3.2 × 10^6^ | 1.3 × 10^6^ | 5.4 × 10^6^ | 1.4 × 10^6^ | 5.2 × 10^6^ | 3.8 × 10^6^ | 2.9 × 10^6^ | 1.7 × 10^6^ | 5.1 × 10^6^ |
|  | 4 | 1.5 × 10^6^ | 2.5 × 10^6^ | 2.7 × 10^6^ | 9.1 × 10^5^ | 2.9 × 10^6^ | 7.7 × 10^5^ | 1.1 × 10^6^ | 1.3 × 10^6^ | 2.9 × 10^6^ |
| Microaerobic | 0 | 3.2 × 10^6^ | 2.4 × 10^6^ | 2.3 × 10^6^ | 2.7 × 10^6^ | 3.8 × 10^6^ | 2.8 × 10^6^ | 2.1 × 10^6^ | 1.7 × 10^6^ | 4.5 × 10^6^ |
|  | 4 | 2.3 × 10^5^ | 2.4 × 10^6^ | 1.2 × 10^6^ | 6.7 × 10^5^ | 2.4 × 10^6^ | 5.9 × 10^5^ | 5.7 × 10^5^ | 7.2 × 10^5^ | 3.7 × 10^5^ |
|  | 8 | 1.2 × 10^5^ | 2.1 × 10^6^ | 7.2 × 10^5^ | 2.6 × 10^5^ | 6.5 × 10^5^ | 2.5 × 10^5^ | 3.4 × 10^5^ | 2.1 × 10^5^ | 2.1 × 10^5^ |
| Anaerobic* | 0 | 3.7 × 10^6^ | 1.9 × 10^6^ | 5.7 × 10^6^ | 1.3 × 10^6^ | 5.2 × 10^6^ | 3.5 × 10^6^ | 2.3 × 10^6^ | 1.5 × 10^6^ | 5.6 × 10^6^ |
|  | 4 | 5.1 × 10^4^ | 2.3 × 10^4^ | 6.1 × 10^4^ | 6.9 × 10^4^ | 2.9 × 10^4^ | 2.5 × 10^4^ | 5.9 × 10^3^ | 6.8 × 10^4^ | 3.7 × 10^4^ |
| **Culture medium used: ASW_Pectin** | | | | | | | | | | |
| Aerobic | 0 | 3.5 × 10^6^ | 1.8 × 10^6^ | 5.4 × 10^6^ | 1.4 × 10^6^ | 5.2 × 10^6^ | 3.5 × 10^6^ | 2.9 × 10^6^ | 1.7 × 10^6^ | 5.1 × 10^6^ |
|  | 4 | 5.3 × 10^7^ | 2.5 × 10^8^ | 7.9 × 10^7^ | 1.3 × 10^6^ | 6.5 × 10^7^ | 7.3 × 10^5^ | 1.2 × 10^8^ | 2.1 × 10^5^ | 6.9 × 10^7^ |
| Microaerobic | 0 | 3.7 × 10^6^ | 1.7 × 10^6^ | 2.3 × 10^6^ | 2.3 × 10^6^ | 3.6 × 10^6^ | 2.7 × 10^6^ | 2.5 × 10^6^ | 1.5 × 10^6^ | 4.7 × 10^6^ |
|  | 4 | 8.1 × 10^6^ | 4.5 × 10^7^ | 5.2 × 10^6^ | 2.4 × 10^6^ | 6.8 × 10^6^ | 2.1 × 10^6^ | 3.7 × 10^7^ | 4.1 × 10^5^ | 9.1 × 10^6^ |
|  | 8 | 2.7 × 10^7^ | 8.9 × 10^7^ | 1.8 × 10^7^ | 1.2 × 10^6^ | 2.7 × 10^7^ | 6.0 × 10^5^ | 8.9 × 10^7^ | 1.1 × 10^5^ | 2.1 × 10^7^ |
| Anaerobic* | 0 | 3.1 × 10^6^ | 1.9 × 10^6^ | 5.1 × 10^6^ | 1.9 × 10^6^ | 4.9 × 10^6^ | 3.5 × 10^6^ | 2.7 × 10^6^ | 1.3 × 10^6^ | 5.5 × 10^6^ |
|  | 4 | 2.9 × 10^4^ | 2.7 × 10^4^ | 8.1 × 10^5^ | 9.1 × 10^4^ | 5.2 × 10^4^ | 3.2 × 10^4^ | 1.9 × 10^4^ | 1.6 × 10^4^ | 3.5 × 10^4^ |
| **Culture medium used: ASW_Xylan** | | | | | | | | | | |
| Aerobic | 0 | 3.5 × 10^6^ | 1.7 × 10^6^ | 5.9 × 10^6^ | 1.4 × 10^6^ | 5.1 × 10^6^ | 3.5 × 10^6^ | 2.9 × 10^6^ | 1.7 × 10^6^ | 5.1 × 10^6^ |
|  | 4 | 6.9 × 10^5^ | 1.1 × 10^6^ | 3.9 × 10^6^ | 1.2 × 10^6^ | 2.9 × 10^6^ | 2.4 × 10^6^ | 1.9 × 10^6^ | 1.1 × 10^6^ | 2.1 × 10^6^ |
| Microaerobic | 0 | 3.5 × 10^6^ | 1.8 × 10^6^ | 2.6 × 10^6^ | 2.9 × 10^6^ | 3.5 × 10^6^ | 2.1 × 10^6^ | 2.5 × 10^6^ | 1.7 × 10^6^ | 4.0 × 10^6^ |
|  | 4 | 3.1 × 10^6^ | 5.7 × 10^5^ | 2.9 × 10^5^ | 8.1 × 10^5^ | 8.7 × 10^5^ | 5.9 × 10^5^ | 5.9 × 10^5^ | 5.8 × 10^5^ | 8.7 × 10^5^ |
|  | 8 | 2.1 × 10^6^ | 4.1 × 10^5^ | 2.1 × 10^5^ | 3.5 × 10^5^ | 2.4 × 10^5^ | 2.3 × 10^5^ | 1.5 × 10^5^ | 1.9 × 10^5^ | 2.5 × 10^5^ |
| Anaerobic* | 0 | 3.8 × 10^6^ | 1.9 × 10^6^ | 5.3 × 10^6^ | 1.3 × 10^6^ | 5.6 × 10^6^ | 3.5 × 10^6^ | 2.2 × 10^6^ | 1.8 × 10^6^ | 5.3 × 10^6^ |
|  | 4 | 1.9 × 10^4^ | 2.2 × 10^5^ | 3.5 × 10^5^ | 5.3 × 10^4^ | 4.8 × 10^3^ | 9.1 × 10^4^ | 2.8 × 10^5^ | 1.3 × 10^4^ | 2.3 × 10^4^ |
| **Culture medium used: ASW_Benzoate** | | | | | | | | | | |
| Aerobic | 0 | 1.8 × 10^6^ | 4.6 × 10^6^ | 4.4 × 10^6^ | 1.2 × 10^6^ | 5.6 × 10^6^ | 1.9 × 10^6^ | 1.6 × 10^6^ | 5.4 × 10^6^ | 1.6 × 10^6^ |
|  | 4 | 5.7 × 10^8^ | 2.8 × 10^7^ | 1.5 × 10^8^ | 2.3 × 10^7^ | 6.8 × 10^8^ | 2.4 × 10^8^ | 1.3 × 10^6^ | 5.6 × 10^6^ | 1.5 × 10^6^ |
| Microaerobic | 0 | 2.6 × 10^6^ | 1.6 × 10^6^ | 3.1 × 10^6^ | 5.7 × 10^6^ | 2.1 × 10^6^ | 1.2 × 10^6^ | 2.6 × 10^6^ | 1.2 × 10^6^ | 4.1 × 10^6^ |
|  | 4 | 1.1 × 10^6^ | 1.4 × 10^6^ | 3.5 × 10^5^ | 1.1 × 10^6^ | 1.2 × 10^6^ | 6.0 × 10^5^ | 5.4 × 10^5^ | 8.5 × 10^5^ | 2.5 × 10^5^ |
|  | 8 | 6.6 × 10^5^ | 2.2 × 10^5^ | 1.1 × 10^5^ | 2.6 × 10^4^ | 1.4 × 10^5^ | 8.4 × 10^5^ | 1.3 × 10^5^ | 4.8 × 10^5^ | 1.2 × 10^5^ |
| Anaerobic* | 0 | 1.6 × 10^6^ | 4.6 × 10^6^ | 4.4 × 10^6^ | 1.2 × 10^6^ | 1.3 × 10^6^ | 1.9 × 10^6^ | 1.8 × 10^6^ | 5.4 × 10^6^ | 1.6 × 10^6^ |
|  | 4 | 1.1 × 10^4^ | 3.5 × 10^4^ | 2.3 × 10^5^ | 7.5 × 10^4^ | 4.5 × 10^3^ | 6.2 × 10^5^ | 5.2 × 10^4^ | 4.2 × 10^4^ | 1.9 × 10^3^ |
| **Culture medium used: ASW_Starch** | | | | | | | | | | |
| Aerobic | 0 | 1.8 × 10^6^ | 1.4 × 10^6^ | 2.5 × 10^6^ | 1.2 × 10^6^ | 1.3 × 10^6^ | 1.9 × 10^6^ | 1.7 × 10^6^ | 3.7 × 10^6^ | 1.1 × 10^6^ |
|  | 4 | 1.6 × 10^6^ | 1.9 × 10^7^ | 7.9 × 10^6^ | 5.8 × 10^5^ | 2.6 × 10^7^ | 4.1× 10^4^ | 3.6 × 10^5^ | 1.1 × 10^7^ | 2.3 × 10^8^ |
| Microaerobic | 0 | 2.6 × 10^6^ | 1.6 × 10^6^ | 3.1 × 10^6^ | 5.7 × 10^6^ | 2.1 × 10^6^ | 1.2 × 10^6^ | 2.6 × 10^6^ | 1.2 × 10^6^ | 4.1 × 10^6^ |
|  | 4 | 6.3 × 10^5^ | 2.4 × 10^6^ | 2.8 × 10^6^ | 2.7 × 10^5^ | 1.7 × 10^7^ | 1.1 × 10^4^ | 1.3 × 10^5^ | 2.2 × 10^6^ | 1.6 × 10^5^ |
|  | 8 | 2.9 × 10^5^ | 3.3 × 10^7^ | 1.9 × 10^6^ | 1.1 × 10^5^ | 4.9 × 10^7^ | 5.3 × 10^3^ | 5.2 × 10^4^ | 8.1 × 10^6^ | 4.5 × 10^4^ |
| Anaerobic* | 0 | 1.6 × 10^6^ | 4.6 × 10^6^ | 4.7 × 10^6^ | 1.2 × 10^6^ | 1.3 × 10^6^ | 1.9 × 10^6^ | 1.8 × 10^6^ | 3.7 × 10^6^ | 4.8 × 10^6^ |
|  | 4 | 3.3 × 10^4^ | 1.2 × 10^4^ | 1.6 × 10^5^ | 4.1 × 10^5^ | 1.2 × 10^3^ | 2.6 × 10^5^ | 3.7 × 10^4^ | 1.2 × 10^4^ | 5.1 × 10^4^ |
|  | 4 | 6.9 × 10^5^ | 1.1 × 10^6^ | 3.9 × 10^6^ | 1.2 × 10^6^ | 2.9 × 10^6^ | 2.4 × 10^6^ | 1.9 × 10^6^ | 1.1 × 10^6^ | 2.1 × 10^6^ |

* Anaerobic incubation involved supplementation of the media with the six terminal electron acceptors NaNO_3_, NaNO_2,_ Fe_2_O_3_, MnO_2_, Na_2_SO_4_, (CH_3_)_2_SO and (CH_3_)_3_NO.

**Table S9.** Increase or decrease in the CFU count of the nine OMZ-sediment isolates upon aerobic, microaerobic, or anaerobic incubation in ASW-based media supplemented with extremely low concentration (1 × 10^-12^ g L^-1^) of yeast extract (ASW_ELY).

| Incubation condition | Days of incubation | *Brevibacterium* sp. BDJS002 | *Brevibacterium* sp. JSBI002 | *Brucella* sp. JSBI001 | *Gordonia hongkongensis*  SMJS1 | *Halomonas* sp. BDJS001 | *Halopseudomonas*  *bauzanensis* SMJS2 | *Marinobacter* *segnicrescens*  AN1 | *Mesobacillus* sp. SBJS01 | *Stenotrophomonas* sp. SBJS02 |
| --- | --- | --- | --- | --- | --- | --- | --- | --- | --- | --- |
| **Initial cell density: 10^6^ cells mL^-1^ medium** | | | | | | | | | | |
| Aerobic | 0 | 1.7 × 10^6^ | 2.3 × 10^6^ | 3.4 × 10^6^ | 3.1 × 10^6^ | 1.2 × 10^6^ | 1.5 × 10^6^ | 5.8 × 10^6^ | 3.3 × 10^6^ | 4.1 × 10^6^ |
|  | 4 | 1.3 × 10^6^ | 9.6 × 10^5^ | 3.2 × 10^6^ | 2.9 × 10^6^ | 9.9 × 10^5^ | 1.4 × 10^6^ | 4.3 × 10^6^ | 2.5 × 10^6^ | 2.7 × 10^6^ |
|  | 8 | 9 × 10^5^ | 2.9 × 10^5^ | 2.4 × 10^6^ | 1.1 × 10^6^ | 5.3 × 10^5^ | 7.8 × 10^5^ | 1.1 × 10^6^ | 1.8 × 10^6^ | 5.9 × 10^5^ |
|  | 28 | 1.1 × 10^5^ | 1.3 × 10^5^ | 1.2 × 10^6^ | 5.2 × 10^5^ | 2.2 × 10^5^ | 1.9 × 10^5^ | 7.1 × 10^5^ | 3.9 × 10^5^ | 5.1 × 10^5^ |
| Microaerobic | 0 | 4.5 × 10^6^ | 3.1 × 10^6^ | 1.5 × 10^6^ | 1.2 × 10^6^ | 1.7 × 10^6^ | 2.6 × 10^6^ | 2.5 × 10^6^ | 5.9 × 10^6^ | 2.8 × 10^6^ |
|  | 4 | 2.7 × 10^6^ | 5.7 × 10^5^ | 1.1 × 10^6^ | 7.7 × 10^5^ | 2.9 × 10^5^ | 3.7 × 10^5^ | 5.7 × 10^5^ | 4.7 × 10^6^ | 2.7 × 10^5^ |
|  | 8 | 1.6 × 10^6^ | 4.8 × 10^5^ | 8.7 × 10^5^ | 2.5 × 10^5^ | 2.1 × 10^5^ | 2.7 × 10^5^ | 3.7 × 10^5^ | 2.9 × 10^6^ | 1.8 × 10^5^ |
|  | 28 | 5.3 × 10^4^ | 2.1 × 10^4^ | 1.9 × 10^5^ | 4.9 × 10^4^ | 6.7 × 10^4^ | 6.1 × 10^4^ | 2.5 × 10^5^ | 2.1 × 10^5^ | 4.2 × 10^4^ |
| Anaerobic | 0 | 4.1 × 10^6^ | 1.4 × 10^6^ | 2.5 × 10^6^ | 3.1 × 10^6^ | 4.0 × 10^6^ | 1.5 × 10^6^ | 2.3 × 10^6^ | 3.1 × 10^6^ | 1.2 × 10^6^ |
|  | 4 | 1.6 × 10^6^ | 3.9 × 10^3^ | 1.2 × 10^6^ | 1.4 × 10^4^ | 1.5 × 10^4^ | 5.1 × 10^4^ | 1.5 × 10^3^ | 1.9 × 10^6^ | 8.8 × 10^3^ |
|  | 8 | 1.3 × 10^4^ | 1.7 × 10^3^ | 1.1 × 10^6^ | 3.9 × 10^3^ | 2.8 × 10^3^ | 2.9 × 10^3^ | 5.7 × 10^2^ | 2.9 × 10^5^ | 3.7 × 10^3^ |
|  | 28 | 0 | 0 | 0 | 0 | 0 | 0 | 0 | 0 | 0 |
| **Initial cell density: 10^3^ cells mL^-1^ medium** | | | | | | | | | | |
| Aerobic | 0 | 1.3 × 10^3^ | 4.1 × 10^3^ | 2.6 × 10^3^ | 3.2 × 10^3^ | 2.8 × 10^3^ | 2.3 × 10^3^ | 3.9 × 10^3^ | 1.0 × 10^3^ | 4.0 × 10^3^ |
|  | 4 | 1.2 × 10^5^ | 5.1 × 10^5^ | 8.2 × 10^4^ | 1.9 × 10^4^ | 5.2 × 10^4^ | 2.4 × 10^5^ | 1.7 × 10^5^ | 1.2 × 10^5^ | 5.0 × 10^4^ |
|  | 8 | 1.1 × 10^5^ | 5.2 × 10^5^ | 8.1 × 10^4^ | 1.8 × 10^4^ | 4.9 × 10^4^ | 2.2 × 10^5^ | 1.8 × 10^5^ | 1.1 × 10^5^ | 4.8 × 10^4^ |
|  | 28 | 8.2 × 10^4^ | 3.5 × 10^5^ | 7.6 × 10^4^ | 1.6 × 10^4^ | 2.8 × 10^4^ | 1.6 × 10^5^ | 1.5 × 10^5^ | 9.3 × 10^4^ | 3.2 × 10^4^ |
| Microaerobic | 0 | 6.7 × 10^3^ | 2.1 × 10^3^ | 3.1 × 10^3^ | 1.3 × 10^3^ | 2.7 × 10^3^ | 5.5 × 10^3^ | 1.5 × 10^3^ | 3.7 × 10^3^ | 2.1 × 10^3^ |
|  | 4 | 8.5 × 10^4^ | 5.2 × 10^4^ | 4.9 × 10^4^ | 6.1 × 10^3^ | 1.6 × 10^4^ | 2.4 × 10^5^ | 2.1 × 10^4^ | 3.3 × 10^4^ | 2.4 × 10^4^ |
|  | 8 | 1.7 × 10^5^ | 1.1 × 10^5^ | 5.6 × 10^4^ | 6.9 × 10^3^ | 3.1 × 10^4^ | 3.7 × 10^5^ | 9.3 × 10^4^ | 8.7 × 10^4^ | 2.5 × 10^4^ |
|  | 28 | 7.2 × 10^4^ | 8.5 × 10^4^ | 1.6 × 10^4^ | 6.1 × 10^3^ | 2.3 × 10^4^ | 2.6 × 10^5^ | 6.5 × 10^4^ | 6.3 × 10^4^ | 1.2 × 10^4^ |
| Anaerobic | 0 | 4.1 × 10^3^ | 3.6 × 10^3^ | 2.9 × 10^3^ | 2.3 × 10^3^ | 2.3 × 10^3^ | 1.7 × 10^3^ | 4.7 × 10^3^ | 2.8 × 10^3^ | 2.3 × 10^3^ |
|  | 4 | 3.8 × 10^3^ | 7.8 × 10^2^ | 1.8 × 10^3^ | 1.4 × 10^1^ | 8.1 × 10^1^ | 2.8 × 10^2^ | 1.2 × 10^1^ | 1.0 × 10^3^ | 1.5 × 10^3^ |
|  | 8 | 1.7 × 10^3^ | 2.1 × 10^2^ | 1.3 × 10^3^ | 1.1 × 10^1^ | 3.9 × 10^1^ | 1.3 × 10^2^ | 1.1 × 10^1^ | 5.9 × 10^2^ | 1.2 × 10^3^ |
|  | 28 | 0 | 0 | 0 | 0 | 0 | 0 | 0 | 0 | 0 |
